# Structural studies of thyroid peroxidase show the monomer interacting with autoantibodies in thyroid autoimmune disease

**DOI:** 10.1101/2019.12.15.876789

**Authors:** Daniel E. Williams, Sarah N. Le, David E. Hoke, Peter G. Chandler, Monika Gora, Marlena Godlewska, J. Paul Banga, Ashley M. Buckle

**Affiliations:** Department of Biochemistry and Molecular Biology, Biomedicine Discovery Institute, Monash University, Clayton, Victoria 3800, Australia; Institute of Biochemistry and Biophysics, Polish Academy of Sciences, Warsaw, Poland; Department of Biochemistry and Molecular Biology, Centre of Postgraduate Medical Education, Warsaw, Poland; Emeritus, King’s College London School of, London, United Kingdom

**Keywords:** Autoimmune thyroid disease, thyroid peroxidase, autoantigen, electron microscopy

## Abstract

Thyroid peroxidase (TPO) is a critical membrane-bound enzyme involved in the biosynthesis of multiple thyroid hormones, and is a major autoantigen in autoimmune thyroid diseases such as Graves’ disease and Hashimoto’s thyroiditis. Here we report the biophysical and structural characterisation of two novel TPO constructs containing only the ectodomain of TPO and lacking the propeptide. Both constructs were enzymatically active and able to bind the patient-derived TR1.9 autoantibody. Analytical ultra-centrifugation data suggests that TPO can exist as both a monomer and a dimer. Combined with negative stain electron microscopy and molecular dynamics simulations, these data show that TR1.9 autoantibody preferentially binds the TPO monomer, revealing conformational changes that bring together previously disparate residues into a continuous epitope. In addition to providing plausible structural models of a TPO-autoantibody complex, this study provides validated TPO constructs that will facilitate further characterization, and advances our understanding of the structural, functional and antigenic characteristics of TPO, a molecule behind some of the most common autoimmune diseases.

## Introduction

Thyroid Peroxidase (TPO) is an enzyme in the thyroid gland responsible for oxidising iodide ions to form iodine (mediated by hydrogen peroxide), which can then be incorporated into the key thyroid hormones triiodothyronine (T_3_) and thyroxine (T_4_) ^1^. These hormones are critical in the regulation of metabolism. TPO is of clinical significance, as it is a target of autoantibodies and autoreactive T cells in autoimmune thyroid diseases (AITD) such as destructive thyroiditis (Hashimoto’s disease) and hyperthyroidism (Grave’s disease) ^2, 3^. AITDs are some of the most common autoimmune diseases in the developed world, with Hashimoto’s disease being a strong risk factor for thyroid cancer ^4^. TPO is suspected to be involved in the pathogenesis of Hashimoto’s thyroiditis (prevalence 300-2980 cases per 100,000 in the Western world), leading to thyrocyte destruction via CD8+ T-cell infiltration resulting in hypothyroidism ^1, 3, 5^. This incidence rate can be compared to other autoimmune diseases – for example type 1 diabetes, which has an incidence in the developed world of 310-570 cases per 100,000 patients - demonstrating the immense disease burden caused by AITDs ^5^. The pathogeneses that underlies these autoimmune diseases is complex, however the lack of any tertiary or quaternary structure of TPO complicates matters. The absence of a structure in which to understand binding of anti-TPO antibodies, which are prevalent almost ubiquitously (>95%) in cases of destructive thyroiditis, complicates the understanding of the pathogenesis and nature of the disease. The precise mechanism by which these antibodies cause damage is uncertain ^2^. Additionally, in some cases AITDs can occur without these autoantibodies being present, and transplacental passage of anti-TPO antibodies does not necessarily cause thyroid damage in the offspring ^3, 6, 7^. Despite this, transplacental passage of these autoantibodies can potentially have cognitive effects on the child. In addition, antibodies to thyroglobulin (Tg) and thyroid stimulating hormone receptor have been identified in both conditions, indicating that there may be multiple antigens in AITD pathogenesis ^2^.

TPO is a member of the animal heme peroxidase family, which also includes myeloperoxidase (MPO), eosinophil peroxidase (EPO) and lactoperoxidase (LPO). This family is characterised by high sequence identity and their iron containing heme groups required for their peroxidase activity. Additionally, a conserved calcium binding site (TPO: His261) is observed across the family, which is critical for coordination of the heme group into the active site ^8^. Proteins within this family exist as both monomers (EPO, LPO) and well as dimers (MPO) *in vivo* ^1^. Additionally, TPO is extensively post-translationally modified (PTM) via N- and O-glycans, another feature of the family. TPO however is the only member of this family which is a transmembrane protein ^9, 10^. TPO is a 933 amino acid long, 107 kDa transmembrane glycoprotein made up of several domains: a heme-containing and catalytically active myeloperoxidase (MPO)-like domain, a complement control protein (CCP)-like domain, an epidermal growth factor (EGF)-like domain, a transmembrane domain and an intracellular domain (Figure 1)^1, 11^. The known homologues of TPO, including LPO and MPO, have 48% and 47% sequence similarity respectively (in relation to the MPO-like domain within TPO), and both LPO and MPO are crystallisable with known structures ^12, 13^. However, no known structure of TPO exists despite previously reported crystals, due to poor diffraction ^14, 15^. The MPO-like domain contains the active site with the catalytically important heme group that is covalently attached to Glu408 and Asp260, as well as His261 necessary for calcium binding ^1^. This domain is highly alpha helical. Both the CCP and EGF-like domains are small domains that are both β-strand rich ^8^.

**Figure 1.**
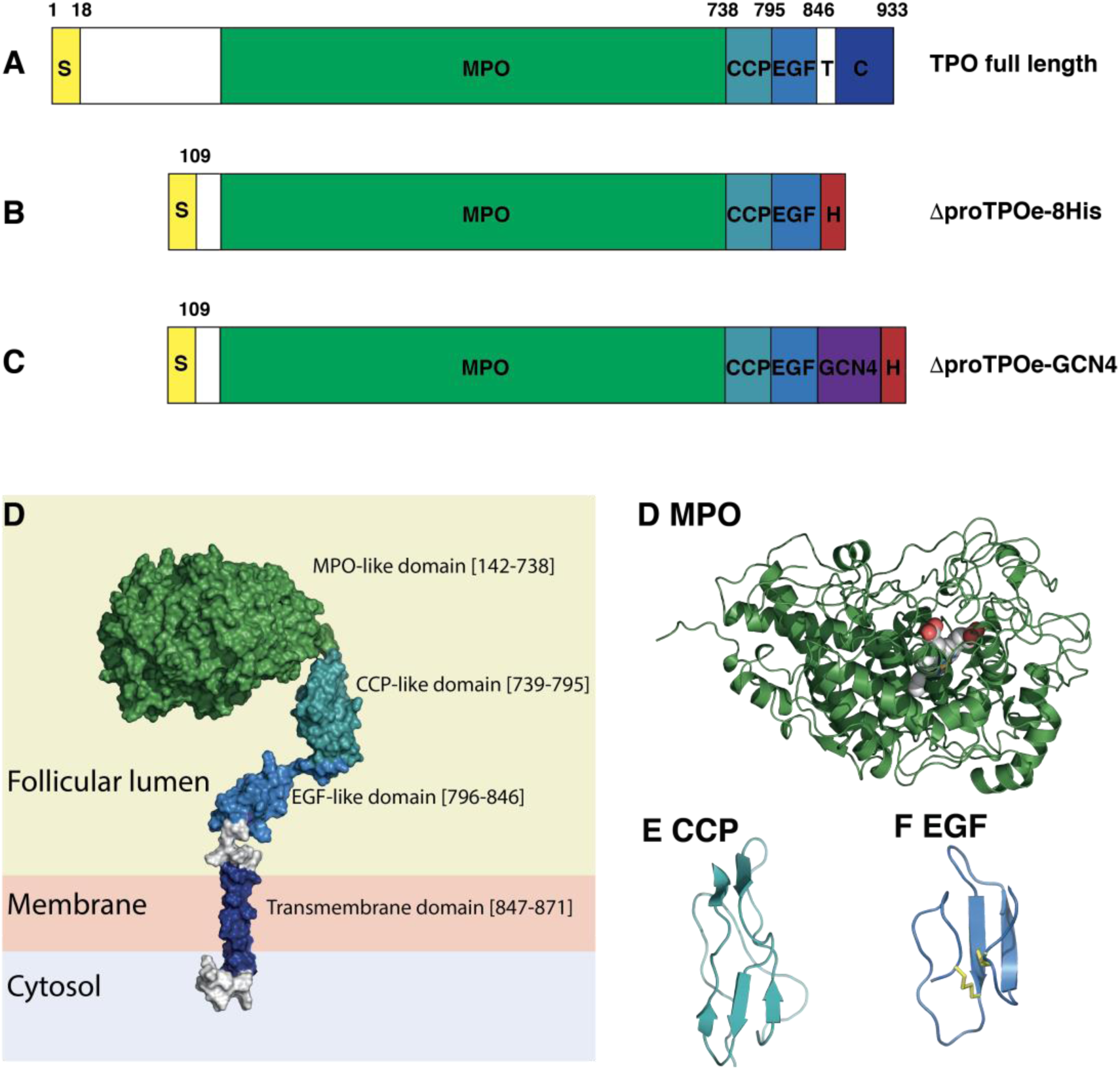
Schematic showing engineered TPO constructs. **(A)** Full length human TPO. **(B)** TPO ectodomain lacking the propeptide containing a C-terminal 8x His tag (ΔproTPOe-8His). **(C)** Construct shown in **(B)** fused to GCN4 (ΔproTPOe-GCN4). S, signal peptide; MPO, MPO-like domain; CCP, CCP-like domain; EGF, EGF-like domain; T, transmembrane span; C, cytoplasmic tail; GCN4, yeast general control protein; H, 8x Histidine tag. **(D)** Schematic showing domain organisation of a TPO monomer in its *trans* configuration. **(E)** X-ray crystal structure of human myeloperoxidase (MPO), with catalytic heme group shown as spheres (PDB ID: 1CXP) ^13^. **(F)** NMR solution structure of the Vaccinia virus complement control protein (PDB ID: 1VVD) ^52^. **(G)** NMR solution structure of a covalently linked pair of EGF-like domains from human fibrillin-1 (PDB ID: 1EMO) ^53^. Disulfide linkages are shown as yellow sticks.

To date, the question of whether TPO exists as a monomer or as a dimer remains open. MPO functions as a dimer, and the conserved cysteine at residue 296 in TPO’s MPO-like domain would suggest a likely site of dimerisation. Despite this, no evidence revealing where in the cell TPO dimers are formed is available, however it has been shown that MPO dimerises after leaving the ER ^10, 16^. In contrast, it has been shown that MPO can exist and function as a monomer *in vivo* despite usually presenting as a dimer ^16^. In the absence of structural characterisation, we previously modelled TPO as homodimer bound by the conserved cysteine residue Cys296 ^8^. Due to the location of this linkage, symmetry restraints and evidence from the primary sequence, it was proposed that the dimer can be modelled in two plausible ways – with the active site of the MPO-like domain facing toward (*cis*) or away (*trans*) from the thyrocyte membrane.

Epitope mapping studies using patient derived autoantibodies against TPO have revealed a pattern of antibody recognition sites in two distinct regions, named immunodominant region A (IDR-A) and immunodominant region B (IDR-B) (Table S1) ^10^. Mapping these regions onto models of a TPO dimer showed that the MPO-like domain was the dominant locale of both IDRs ^8^. IDR-A was not closely clustered on either the *cis* or *trans* model, and the epitopes in some cases were too sparse to be engaged by a single antibody. This would suggest that TPO may exhibit flexibility allowing these regions to coalesce into one discrete epitope. In IDR-B, the epitopes cluster close to the MPO-like domain dimer interface in both *cis* and *trans* models, burying parts of the epitope in this interface. As such, the question of whether TPO exists as a monomer or dimer is central to understanding its autoantigenicity ^1, 8^. Thus, probing the structural characteristics of TPO may provide key insights into the molecular basis of autoimmune disease.

In order to understand whether TPO exists as a monomer or dimer and to provide more insights into its structure, here we report a structural, functional and biophysical characterization of two TPO constructs; ΔproTPOe-8His and ΔproTPOe-GCN4, the latter with a leucine zipper dimerisation motif engineered to stabilise the dimer form.

## Materials and Methods

### Construction of pcDNA/FRT/ΔproTPOe containing a C-terminal 8x His tag

An inverse PCR reaction was performed on pcDNA/FRT5/ΔproTPOe/8His plasmid ^17^ using the following primers: forward 5’-TGATAGTCTAGAGTCGAC-3’ and reverse 5’-GTGGTGGTGGTGGTGGTGGTGGTGAGTCGCCCGAGGGAGCCT-3’ to introduce a C-terminal 8x His tag into the cDNA of the clone. A *DpnI* digestion was performed on the PCR products followed by a blunt end ligation. Ligated products were transformed into chemically competent DH5α cells. Successful colonies were screened and sequenced on both strands.

### Construction of ΔproTPOe containing the yeast GCN4 dimerisation motif

The gene for the yeast GCN4 dimerisation motif was chemically synthesised (GenScript) and subcloned with *NotI and BamHI* into the pUC57 ampicillin resistant vector containing the C-terminal ΔproTPOe sequence. The pcDNA5/FRT/ΔproTPOe and pUC57/GCN4 plasmid was simultaneously digested with *NotI* and *BamHI* and the final products isolated on a 1% agarose gel and subjected to gel purification. Cloning of the yeast GCN4 dimerisation motif into the pcDNA5/FRT/ΔproTPOe/8His plasmid product was achieved by utilising an internal *BamHI* restriction site within the ΔproTPOe protein sequence and a *NotI* site in the vector. Both plasmid products were digested with *BamHI*, ligated and then transformed into chemically competent DH5α cells. Successful colonies were screened and sequenced on both strands.

### Expression in HEK293 cells

EXPI293 cells (ThermoFisher Scientific) were transiently transfected with the ΔproTPOe-8His and ΔproTPOe-GCN4 pcDNA5/FRT plasmids respectively. These cells were diluted to 0.5×10_5_ cells per millilitre 48 hours prior to transfection with FreeStyle 293 Expression Medium (ThermoFisher Scientific). Plasmid DNA isolated above was added to pre-warmed PBS at a ratio of 1 µg/mL of cell culture, in addition to polyethylenimine at a rate of 4µg per 1µg of DNA. Cells were counted and adjusted to 1.5×10_6_ cells/mL which had >95% viability for transfection. This buffer was added to the culture to an amount equal to 10% of the final volume. Glucose concentration was adjusted to 33 mmol/L, and the cells were incubated at 37°C with 5% CO_2_ for 7 days. Final concentrations of 20 µM hematin and 10 µM hydrogen peroxide were added to the culture twice during the course of expression.

### Protein Purification of ΔproTPOe-8His and ΔproTPOe-GCN4 constructs

EXPI293 supernatants of each construct underwent centrifugation at 6,000x g for 5 min to remove any cell debris. ΔproTPOe was purified from conditioned media via immobilised metal affinity chromatography, using a nickel-nitrilotriacetic acid (Ni-NTA, Qiagen) sepharose. Once bound, the resin was washed with wash buffer (PBST-0.05% Tween 20, 10 mM imidazole) to remove loosely and non-specifically bound proteins. ΔproTPOe constructs were eluted with elution buffer (PBST-0.05%, 240 mM imidazole), pooled and concentrated. As a final polishing step, ΔproTPOe was further purified by a HiLoad Superdex S200 16/60 column (GE Healthcare) using gel filtration buffer (1x PBS pH 7.4, 0.01% Tween 20). ΔproTPOe eluted as a single symmetric peak. Protein purity was analysed by SDS-PAGE and Western blotting.

### TR1.9 protein transformation, expression and purification

The TR1.9 Fab sequence contained within a pBP101 plasmid was transformed, expressed and purified as previously published ^18^.

### Enzyme-linked immunosorbent assay (ELISA) of TPO-Fab interaction

An ELISA was performed to determine whether TR1.9 Fab would bind to both TPO constructs, ΔproTPOe-8His and ΔproTPOe-GCN4. The experiment included a number of controls (IgG, positive control, conserpin ^19^, negative control, as well as PBS blanks).

A 96-well polystyrene ELISA plate (Corning) was coated with diluted antigen in PBS (either TPO or control at 10 μg/mL) and incubated at room temperature for 4 hours. The plates were then washed with PBST-0.05 (1x PBS pH 7.4, 0.05% Tween 20). four times and then blocked with 5% skim milk in PBST-0.05, pH 7.4 overnight at 4 °C. The plates were then once again washed four times with PBST-0.05, pH 7.4, after which primary antigen (TR1.9 Fab, 1 mg/mL) was added to the wells at a 1 in 500 dilution in PBST-0.05, and then incubated at room temperature for 30 minutes. The plates were once again washed under the same conditions as stated earlier, before 1:2000 dilutions of the secondary antibodies in PBST-0.05 (anti-human IgG conjugated with horseradish peroxidase (HRP), or anti-His, ThermoFIsher Scientific) were added, followed by another 30-minute incubation at room temperature. After a final wash, detection was performed using a 1-Step Ultra-TMB (*3,3’,5,5’-tetramethylbenzidine)* ELISA solution (ThermoFisher Scientific). A final addition of an equal amount of 100% acetic acid after one hour acted as a stop solution. The absorbance readings were then measured via an endpoint protocol on a BioRad 96-well plate reader, at a wavelength of 450 nm.

### Analysis of ΔproTPOe-TR1.9 Fab complex using size-exclusion chromatography

ΔproTPOe-8His and ΔproTPOe-GCN4 was incubated in two-fold molar excess of TR1.9 Fab, in 1x PBS, pH 7.4 supplemented with + 0.01% Tween 20. This complex was incubated at 4 *°*C for 30 minutes before injection onto a Superdex S200 16/60 size-exclusion column (GE Healthcare). The sample was run at 1.0 mL/min and eluted in 1.5 mL fractions, before being analysed by SDS-PAGE.

### Western blot analysis

Following SDS-PAGE analysis, the proteins on the gel were transferred to a PVDF membrane at 100V for one hour. The membrane was then blocked in 5% skim milk in PBST-0.05 at room temperature for one hour with shaking. Constructs containing a His tag were detected by a single step horse radish peroxidase (HRP)-conjugated anti-His antibody produced in mouse (ThermoFisher Scientific) at a 1:1000 dilution. TR1.9 Fab was detected by a HRP-conjugated anti-human IgG (Fab specific) at the same dilution for one hour. Both were washed with TBS-T four times before application of an electrochemiluminescence (ECL) solution (GE Healthcare) and visualised by X-ray photographic film (FujiFilm) at various exposure times.

### Mass Spectrometry analysis of ΔproTPOe-GCN4

Samples of ΔproTPOe-GCN4 were run on an SDS-PAGE gel with fresh tricine running buffer and fresh Coomassie brilliant blue stain G-250. Bands were removed from the gel and mass spectrometric analysis using LC-MS/MS was performed at the Monash Biomedical Proteomics Facility. Peptide fragments were compared against a reference ΔproTPOe-GCN4 sequence.

### Guaiacol activity assay

TPO can also oxidise the compound guaiacol, which remains colourless in solution, to tetraguaiacol, which appears orange. This characteristic can be exploited by spectroscopic analysis to record the enzymatic activity of TPO. This reaction was undertaken in 96-well plates, with each reaction containing 0.1 mg/mL TPO sample, 1 mM hydrogen peroxide, 33 mM guaiacol and TPO. Detection via a Bio-Rad 96-well plate reader occurred after a 5-minute incubation for these reagents, at 450 nm. We also recorded this interaction over time, comparing with a HRP-conjugated antibody (ThermoFisher Scientific, 1 in 1000 dilution) and a PBS blank for controls, recording optical density over time.

### Soret peak analysis to determine incorporation of heme group

The successful incorporation of an iron containing heme group in proteins can be detected by what is known as a Soret peak. ΔproTPOe-8His and ΔproTPOe-GCN4 at a concentration of 0.5 mg/mL in 1x PBS pH 7.4 were placed in a cuvette and a fluorescence spectrum was obtained, with an incident wavelength of 330 nm. A distinctive peak at 385 nm is a Soret peak and is distinctive of hemoproteins. This absorbance spectrum was obtained on a HoribaJobinYvon FluoroMax4 spectrophotometer.

### Protein stability measurements

TPO constructs at a final concentration of 0.3 mg/mL were placed in different buffer conditions ranging from pH 4.0 to 8.0 (Table S2). Unfolding was measured via intrinsic tryptophan fluorescence by heating from 20 to 95 *°*C, with a ramp rate of 1 *°*C/min, using a Prometheus NT.48 instrument (NanoTemper).

### Binding affinity measurements

Bio-layer interferometry experimental data was collected via a BLItz instrument (FortéBio). Ni-NTA biosensors were equilibrated overnight in 1x PBS, 0.01% Tween 20, 1% BSA, pH 7.4 at 4 °C before use. This assay consisted of five steps: initial baseline (30 s), protein loading (150 s), baseline (30 s), association (240 s) and dissociation (480 s). All steps occurred using the same buffer composition listed above. ΔproTPOe-8His was loaded onto the biosensors during the loading phase at a concentration of 50 μg/mL as per the manufacturer’s specifications, with a loading signal of approximately 1 nm. TR1.9 Fab was incubated with the biosensors at concentrations of 0, 50, 100, and 500 nM during the association phase. A blank control reading was used as a baseline during data processing. Using the BLItz Pro version 1.2.1.3 software, the dissociation constant K_D_ (nM), and the rate constants k_a_ (1/Ms) and k_d_ (1/s) were calculated using a 1:1 binding model with global fitting. Curves were adjusted within the BLItz Pro software to match at the start of both association and dissociation in order to adjust for changes in conditions between steps. R_2_ values for the calculated fit were reported as 0.97.

### Analytical ultra-centrifugation analysis (AUC)

AUC can be used to observe the oligomerisation status of proteins, as well as analyse complex formation. ΔproTPOe-8His and ΔproTPOe-GCN4, as well as each in complex with TR1.9 Fab, were ultra-centrifuged in a BeckmanCoulter Analytical Centrifuge using a double sector cell with quartz windows in 4-hole An60-Ti rotors. The wavelengths for further analysis were selected by an initial scan at 3000 RPM at room temperature to select the best radial settings. For the sedimentation velocity experiment, 420 µL of sample and 400 µL of buffer (1x PBS pH 7.4) were placed in each cell and the experiment was run with the radial settings discussed earlier at 40,000 RPM. 500 scans were performed with a rate of 15 scans per minute. Information about buffer viscosity, density and partial volume were determined by the software SEDNTERP. A c(s) sedimentation distribution model was determined using the SEDFIT software available from https://sedfitsedphat.nibib.nih.gov/software/default.aspx.

### Negative stain electron microscopy (EM) analysis

A TPO-Fab complex was formed by addition of a 2x molar excess of TR1.9 Fab to freshly purified ΔproTPOe-8His. This was incubated on ice for 30 min before separation using an analytical S200 10/300 column in 1x PBS pH 7.4 (GE Healthcare). 10 µL of ΔproTPOe-8His alone and in complex with TR1.9 Fab at 0.05 mg/mL were pipetted onto freshly glow discharged carbon grids and stained with 2 % uranyl acetate. Micrographs at 67000x magnification were recorded on a FEI Tecnai Spirit T12 transmission electron microscope (TEM), with a spot size of 1, dose of 20 e/Ås^-1^, and an exposure time of one second. Fitting of the models within the density was performed using the inbuilt commands within UCSF Chimera 1.13.1 ^20^.

### Computational resources

Parametrisation and molecular dynamics (MD) simulations of TPO constructs were performed on in-house hardware using NVIDIA 1080 Ti Pascal GPUs.

### MD systems preparation

Starting structures for *cis* and *trans* variants of ΔproTPOe were derived from the models presented in Le and co-workers ^8^. An extended variant of ΔproTPOe was developed which did not bias the orientation of the CCP-like and EGF-like in relation to the MPO-like domain. Additionally, each structure had separate runs with unrestrained, as well as positionally restrained IDR residues. These residues were weakly constrained with harmonic position restrains of 2 kcal Å^2^ mol^−1^. These proteins contained protonation states appropriate for pH 7.0 (as determined by PROPKA) and were inserted in a rectangular box with a border with a minimum of 12 Å ^21, 22^. They were then explicitly solvated with TIP3P water, sodium counter-ions were added and then the system was parameterised using the AMBER ff99SB forcefield ^23–26^. After 10000 steps of energy minimisation, an equilibration stage was performed. The temperature was raised from 0 K to 300 K with a constant volume and a 10 K ramp over 1 ns, with Langevin temperature coupling relaxation times of 0.5 ps. After the target temperature was achieved, the pressure was equilibrated to 1 atm using the Berendsen algorithm over 0.5 ps ^27^. The MD simulations used periodic boundary conditions and a time step of 2 fs, with temperature maintained at 300 K using the Langevin thermostat and pressure maintained at 1 atm using the Berendsen method as described above. All MD simulations were run in triplicate, with the same starting structure but with altered starting velocities following the equilibration and parametrisation steps. Each run extended for 400 ns using NAMD 2.9 ^28^.

### MD analysis

Manipulation and analysis of the simulations were performed using VMD 1.9.3, MDTraj and custom scripts ^29, 30^. Models of ΔproTPOe were analysed for root mean square deviation (RMSD), with the RMSD of backbone heavy atoms in relation to the initial structure calculated for every 0.1 ns of simulation after calculating a least-square fit. Output structures for further analysis were selected from a plateau in the RMSD calculation, whereby the structure adopted a stable state. Distance between selected IDR residues in each frame was also calculated using VMD and custom scripts. RMSD is given as an average value per residue per 0.1 ns of simulation time. All structural representations were prepared using PyMOL 2.3.2.

## Results

### Expression and Purification of TPO constructs

TPO’s domain structure, the engineered constructs and structural models are shown in Figure 1. The ΔproTPOe-8His construct lacks a propeptide domain, which has previously been shown to not influence TPO secretion, activity or immunogenicity ^17^. Additionally, the expression of only the ectodomain allows ΔproTPOe-8His to be secreted into the media rather than be retained in the membrane. ΔproTPOe-8His was expressed into the media successfully in EXPI293 cells (Figure 2). Purification of the secreted media shows the presence of a major band ∼110 kDa that is ΔproTPOe-8His. Upon a two-step purification involving Ni-NTA affinity chromatography and size-exclusion on a Superdex S200 16/60 column, ΔproTPOe-8His elutes as a single symmetrical peak at 75.5 mL which is consistent with that of a 110 kDa according to our calibration (Figure 2C). There appears to be no other high molecular weight species on the chromatogram. Despite this, the fractions making up this peak when analysed via reducing SDS-PAGE (Figure 2A) demonstrate a number of minor species, which may be due the heterogeneous nature of the glycosylation of TPO or protein degradation from proteases released from cells during the 7-day expression cycle.

**Figure 2.**
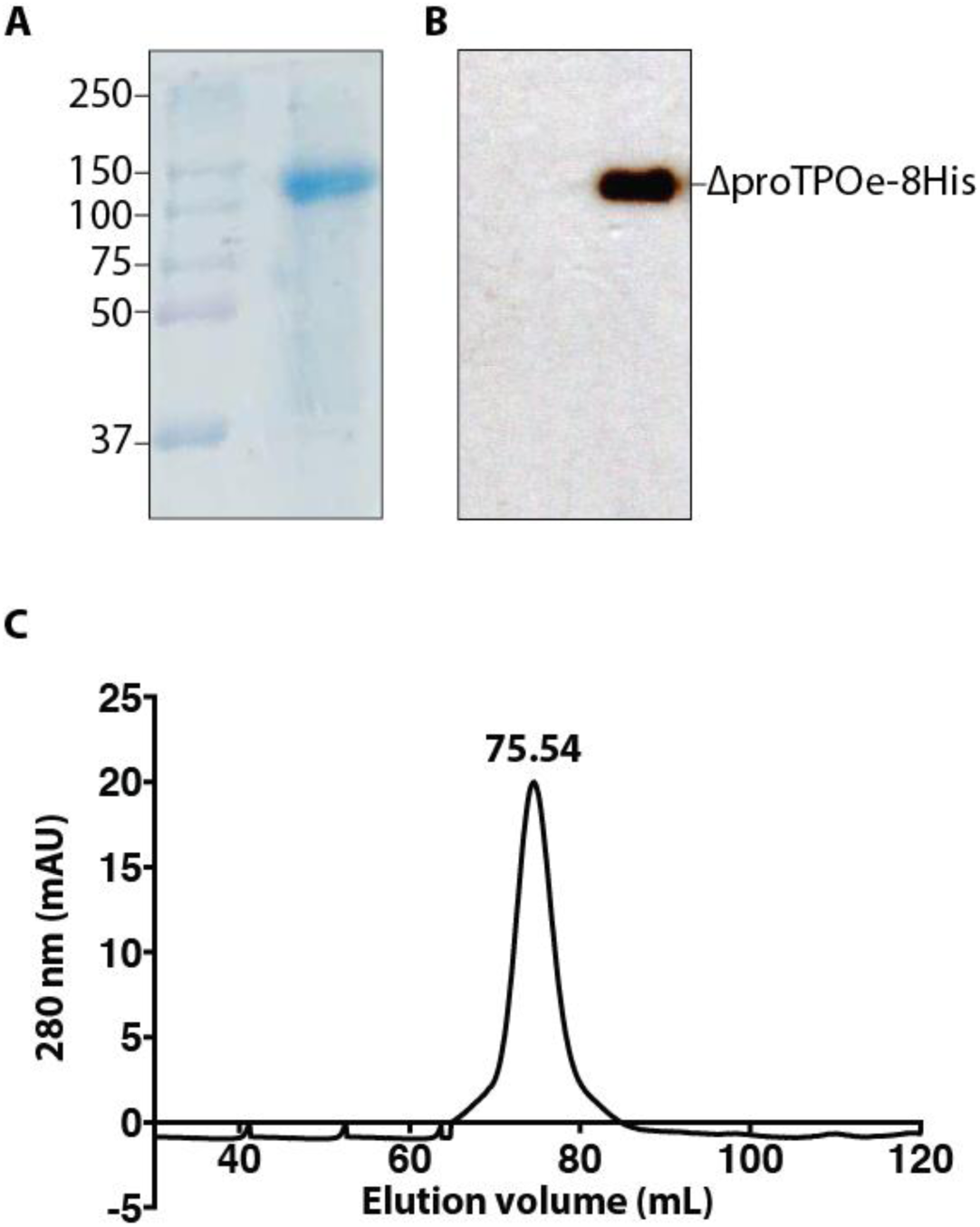
Purification of TPO construct ΔproTPOe-8His. **(A)** SDS-PAGE analysis of purified ΔproTPOe-8His shows a major band at ∼110 kDa. **(B)** Western blot of purified ΔproTPOe-8His with anti-His antibody shows an immunoreactive protein at the same molecular weight as in (A). **(C)** A chromatogram from Superdex S200 16/60 column, showing ΔproTPOe-8His being purified as a single major peak at 75.54 mL.

In an attempt to favour TPO dimerisation we engineered a GCN4 leucine zipper dimerisation motif into the C-terminus of TPO, creating ΔproTPOe-GCN4. Expression and purification proceeded as with the ΔproTPOe-8His construct (Figure S1). SDS-PAGE followed by mass spectroscopic analysis confirmed the presence of intact ΔproTPOe-GCN4 (∼110 kDa) (Figure S2). Despite the quality of our preparations, TPO suffered from a shorter shelf life and often appeared to degrade into a 75kDa component according to SDS-PAGE. This fragment was analysed by mass spectrometry and was identified as a truncated TPO (Figure S3). Enzymatic activity of the constructs was validated by spectroscopic analysis, measuring both proper heme incorporation as well as function via the guaiacol activity assay (Figure 3). Whereas ΔproTPOe-8His is enzymatically active, activity could not be detected for ΔproTPOe-GCN4 (data not shown).

**Figure 3.**
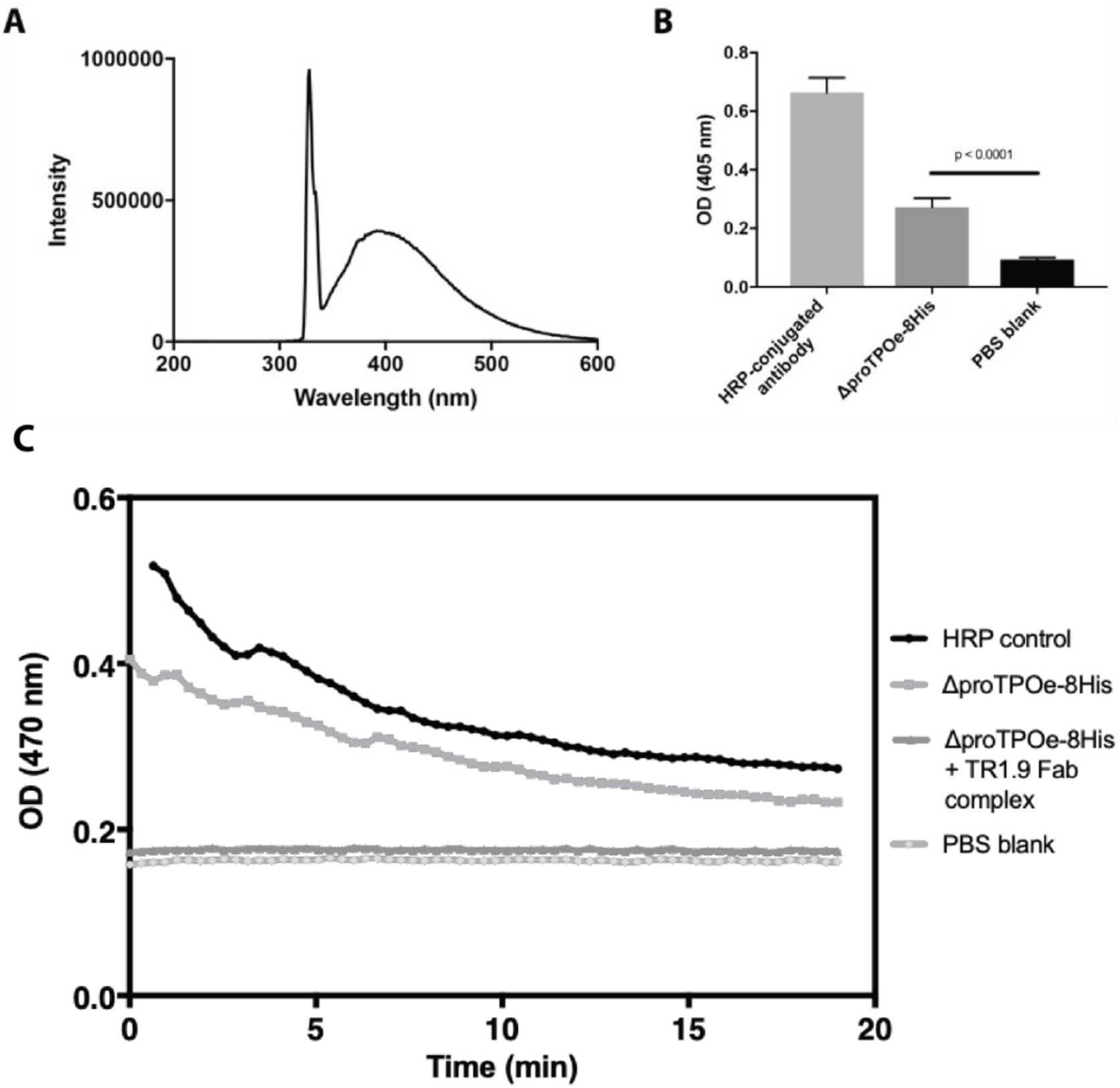
Characterisation of enzyme activity. **(A)** Spectral scan of ΔproTPOe-GCN4. An excitation wavelength of 330nm resulted in a Soret peak at 385nm, which is characteristic of hemoproteins. This indicates successful heme group incorporation into ΔproTPOe-GCN4. Intensity is given in arbitrary units. **(B)** TPO activity as measured by the guaiacol activity assay. Optical density was recorded at 405nm for the three samples after 10 minutes, including an HRP positive control. ΔproTPOe-8His shows statistically significant activity indicating that it is enzymatically active. Error bars are standard deviation from the mean and statistical tests performed with a two-tailed t-test with with a 95% confidence interval. All samples were performed in quadruplicate. **(C)** ΔproTPOe-8His activity is prevented in the presence of TR1.9 Fab during the guaiacol assay.

ΔproTPOe-8His shows statistically significant activity, suggesting that this protein is likely enzymatically active and correctly folded (Figure 3B). Since the thermal stability of TPO has not been reported to date, we measured the stability of the ΔproTPOe-8His construct using thermal unfolding and monitoring intrinsic tryptophan fluorescence in several buffer conditions. The midpoint of unfolding (T_m_) of ΔproTPOe-8His reached a maximum of 55.2 °C at pH 7.0 (Table S2). We next showed that TR1.9 Fab binding completely inhibits the catalytic activity of TPO (Figure 3C).

### Assessment of Oligomerisation State of TPO Constructs

Chromatographic analysis suggests that the TPO constructs behave as monomers, but we suspected that based upon previous work that the TPO molecule is elongated and relatively flexible, and thus we are unable to unambiguously differentiate between monomers and dimers ^1, 8^. From the size-exclusion chromatography data, it is possible that the monomer and dimer of TPO are not separated if they have the same or similar Stokes radii (R_s_), which may be possible given their symmetry. We therefore next used analytical ultracentrifugation and sedimentation velocity experiments to provide equilibrium information about TPO’s shape, size and oligomeric state. For ΔproTPOe-GCN4, two species could be detected, having a standardised weight-average sedimentation coefficient of 3.6 and 5.2 (Figure 4A). The molecular weights reported by c(M) analysis (not shown) in each case was 114 kDa and 201 kDa respectively, with a frictional ratio of 2.36. These molecular weights are consistent with a monomer (∼110 kDa) and a dimer (∼220 kDa) species. The relative abundance of each of the two species was analysed by SEDFIT (using area under the curve), with a monomer:dimer ratio of approximately 1.33:1, indicating the monomer is more abundant. The frictional ratio of 2.36 suggest a non-spherical, elongated shape, consistent with our previous modelling (a frictional ratio of 1 would suggest a perfect sphere) ^8^. For the ΔproTPOe-8His construct (Figure 4B), two species of standardised weight-average sedimentation coefficient of 3.5 and 5.0 were detected, which were similar to the ΔproTPOe-GCN4 construct. The monomer:dimer ratio appears in this case to be closer to 1:1 in terms of distribution. This would indicate that both constructs exist as monomeric and dimeric forms in solution. Calculated Stokes radii are listed in Table S3.

**Figure 4.**
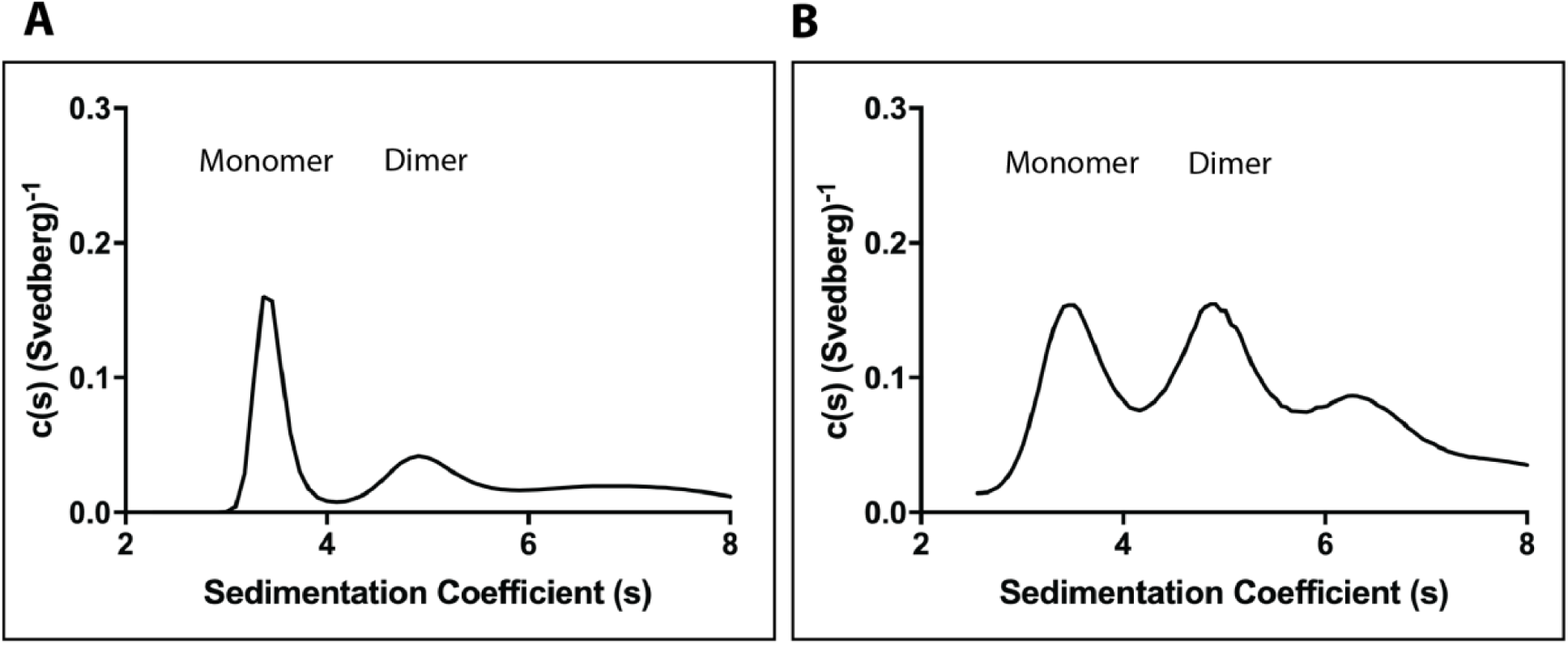
Sedimentation distribution of ΔproTPOe-GCN4 alone and ΔproTPOe-8His alone. **(A)** Two distinct species are detected at a sw(^20,w^) of 3.6 and 5.2 respectively on the sedimentation distribution for the ΔproTPOe-GCN4 construct. (**B)** Two major species are detected at a sw(^20,w^) of 3.5 and 5.0 respectively on the sedimentation distribution in the ΔproTPOe-8His construct.

### Characterisation of a human TPO-autoantibody complex

In order to gain insight into the nature of the interaction between TPO and the patient-derived TR1.9 autoantibody, we expressed the Fab portion of TR1.9 as previously described ^18^. TR1.9 Fab was shown to bind both ΔproTPOe-8His and ΔproTPOe-GCN4 using ELISA (Figure 5). It is apparent from the size-exclusion chromatogram that Fab binds to full-length ΔproTPOe-8His (Figure 5A). There was only a minor peak shift with no appearance of any large species on the chromatogram (dimeric TPO with two Fabs would exceed 320 kDa), which would suggest a stoichiometric monomeric TPO-Fab complex. We also calculated the Stokes radii of the various TPO constructs with and without Fab using analytical size-exclusion chromatography as well as AUC (Table S3). Bio-layer interferometry indicated that ΔproTPOe-8His binds TR1.9 Fab with an affinity of 20 nM (Figure S4). Taken together, this data confirms the antigenic quality of our constructs.

**Figure 5.**
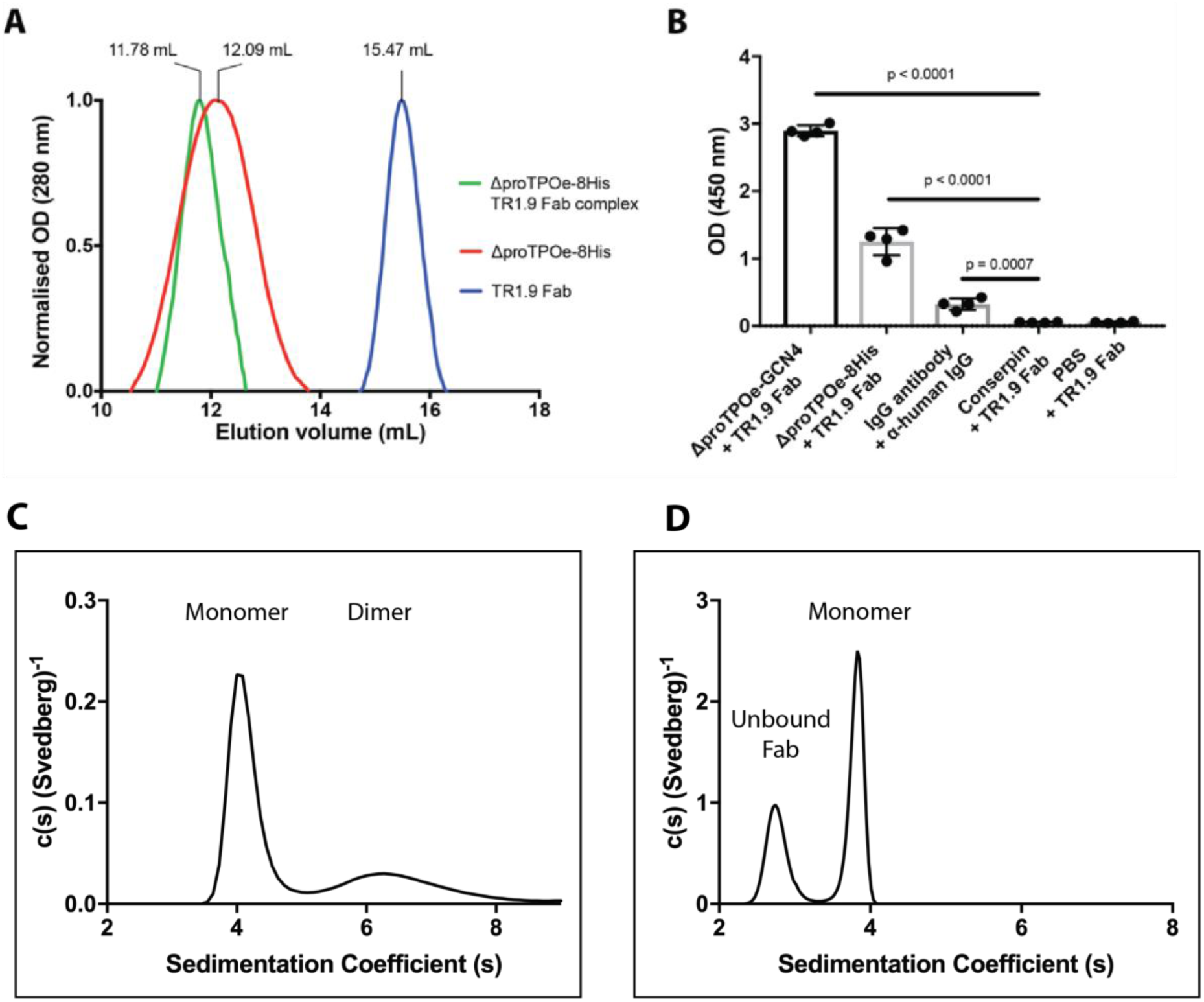
TPO-TR1.9 Fab complex characterization. (**A**) Analytical size-exclusion chromatography of the ΔproTPOe-8His-TR1.9 Fab complex. Elution profiles of ΔproTPOe-8His alone (red), ΔproTPOe-8His-TR1.9 Fab complex (green) and TR1.9 Fab alone (blue). (**B**) ELISA results of TPO-Fab binding. TR1.9 Fab shows statistically significant binding to both TPO constructs, with a p value less than 0.0001 compared to a non-specific protein that does not have the required epitope (conserpin ^19^, negative control), as well as a PBS blank. IgG antibody and anti-human IgG was used as a positive control. Error bars are standard deviation from the mean and statistical tests performed with a two-tailed t-test with a 95% confidence interval. All samples were performed in quadruplicate. **(C)** Sedimentation distribution of ΔproTPOe-GCN4 bound to TR1.9 Fab. Two distinct species are detected at a sw(^20,w^) of 4.1 and 6.6 respectively. **(D)** Sedimentation distribution of ΔproTPOe-8His bound to TR1.9 Fab. Two distinct species are detected at a sw(^20,w^) of 2.8 and 3.9 respectively.

**Figure 6.**
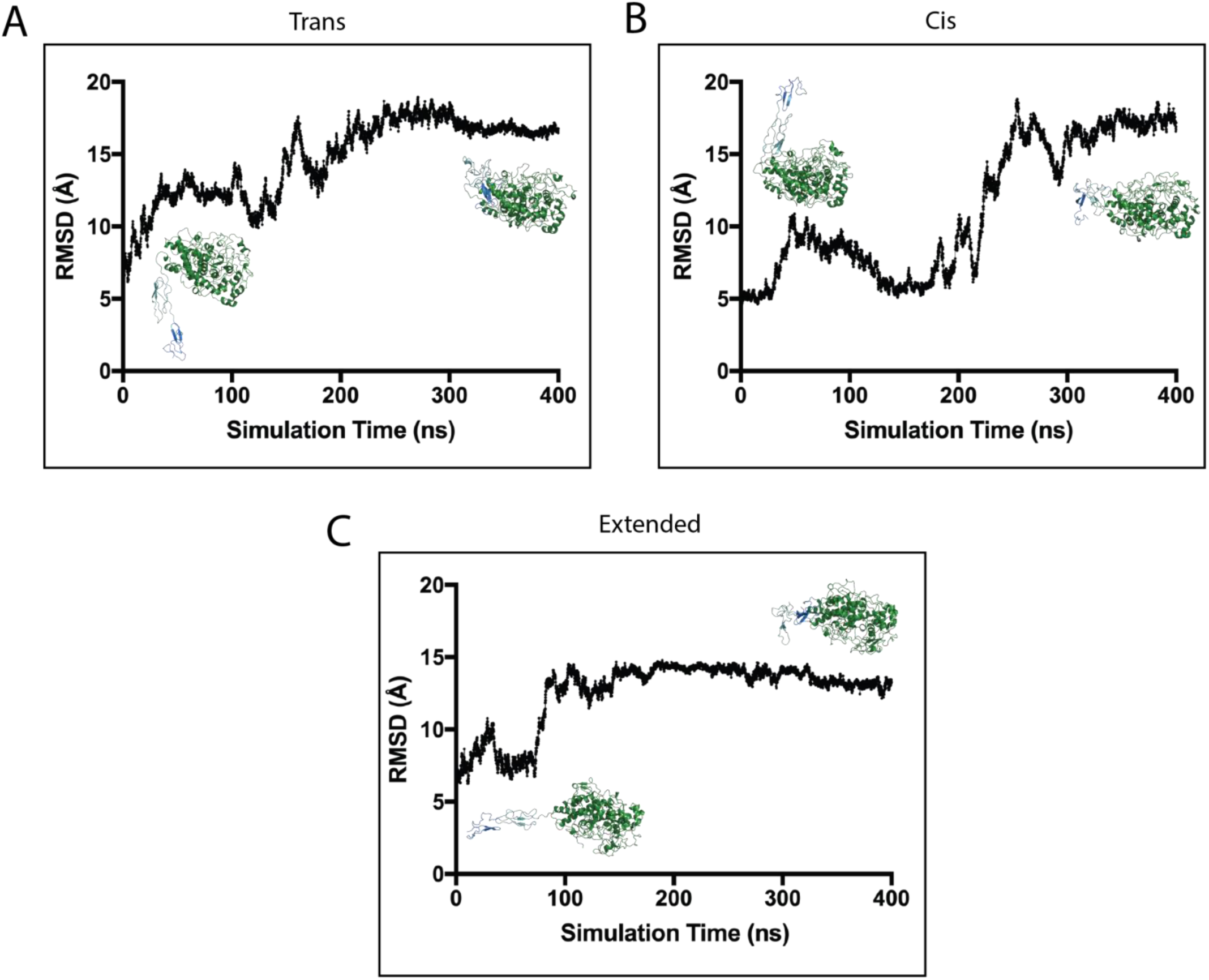
Molecular dynamics simulations of a ΔproTPOe monomer. Root mean square deviations (RMSD) is given as an average value per residue per 0.1ns of simulation time for the (**A)** *trans*, (**B)** *cis* and (**C)** *extended* models of ΔproTPOe. Within each panel is a model representation of a starting structure of the particular form of TPO on the left, with a model taken from the plateau of the MD run on the right-hand side. The MPO-like domain, CCP-like domain and EGF-like domain are coloured in forest green, light teal and marine blue respectively (as in Figure 1).

### Analytical Ultra-centrifugation of the ΔproTPOe-GCN4-TR1.9 Fab Complex

We next used analytical ultra-centrifugation to investigate the association between TR1.9 Fab and ΔproTPOe-GCN4, as well as the effect of TR1.9 Fab binding on the equilibrium of monomer and dimer TPO species. The sedimentation coefficient for TR1.9 Fab alone was determined as sw(^20,w^) = 2.6S. AUC analysis of an equimolar mixture of TR1.9 Fab and ΔproTPOe-GCN4 shows two peaks (Figure 5C). The lack of a peak at 2.6S suggests that none, or very little of the Fab, remained un-complexed. Therefore, the two remaining peaks are most likely the TPO monomer/dimer peaks as observed in the previous experiment (Figure 4A). The observed shift in their standardised weight-average sedimentation coefficients (4.1S, 6.6S, respectively) suggest a change in their shape and mass, indicating Fab binding to both monomer and dimer ΔproTPOe-GCN4. The frictional ratio has also changed to 1.77 (from 2.36 with ΔproTPOe-GCN4 alone), indicating that TPO has taken a more spherical shape upon TR1.9 Fab binding. Importantly, the ratio of monomer and dimer has shifted to approximately 2:1. This indicates that TR1.9 Fab preferentially binds the TPO monomer, thus perturbing the monomer:dimer equilibrium.

### Analytical Ultracentrifugation of ΔproTPOe-8His with TR1.9 Fab

The same sedimentation velocity experiments were undertaken with ΔproTPOe-8His in the presence of TR1.9 Fab, with the Fab in 2-fold excess of TPO. Two distinct species were observed (Figure 5D), with a standardised weight-average sedimentation coefficient of 2.8 and 3.9. This is largely consistent with the Fab alone (2.6S) and ΔproTPOe-GCN4 monomer plus TR1.9 Fab (4.1S) data obtained previously with the alternate construct. Interestingly, in this case there appears to be no other large species, indicating that the binding of Fab has pushed the monomer:dimer equilibrium such that all ΔproTPOe-8His exists exclusively in its monomeric form. AUC further shows that TR1.9 Fab can successfully bind both TPO constructs.

The monomeric ΔproTPOe-8His with Fab has a frictional ratio of 1.90 and a Stokes radius of 57.4, similar to that of the monomeric form of ΔproTPOe-GCN4 bound with Fab (Table S3). Taken together, these data indicate that TPO adopts an elongated, non-spherical structure, consistent with our previous model ^8^. However, a decrease in both the frictional ratio and Stokes radius when bound with TR1.9 Fab suggests that ΔproTPOe-8His adopts a more compact, globular shape upon antibody binding.

### Molecular Dynamics Simulations Reveal Monomer Conformations Compatible with Antibody Binding

Given the structural changes indicated by AUC data, we next used molecular dynamics simulations to explore the structure of TPO monomers in molecular detail. Our previous modelling analysis of epitope mapping data suggest that some residues in IDR-A are too far away to be engaged by a single autoantibody ^8^. One such autoantibody, T13, is reported to bind H353-Y363, P377-R386, K713-S720 and Y766-Q775 ^31–34^. The average maximum dimension of an epitope-containing surface is reported at 28 Å (s.d. of 8 Å) ^35^. In contrast, in our previous models of TPO residues implicated in antibody binding are more disperse, with some residue pairs separated by more than 70 Å ^8^. This discrepancy suggests that TPO may undergo significant conformational change upon antibody binding, such that epitope residues are brought into closer proximity, consistent with our AUC analysis. To explore this hypothesis, we performed molecular dynamics simulations of a ΔproTPOe monomer in three starting configurations: the *cis* and *trans* monomer as generated previously ^8^, and an *extended* conformation in which the CCP-like and EGF-like domains of TPO extend out from the MPO-like domain without any bias in its orientation.

After approximately 250 ns of MD simulation time, the CCP-like and EGF-like domains of the *cis* model (containing the epitope Y766-Q775) move towards the MPO-like domain, such that this epitope coalesces around the T13 epitope residues H353-Y363 and R377-R386 (Figures 7, 8 & S6A). As a result, the average residue separation within the epitope decreases from 56 to 40 Å, reaching 21 Å at some points in the simulation. This distance is such that it could reasonably be engaged by a single autoantibody via a continuous epitope, which may suggest a mechanism by which autoantibodies could engage multiple sites on TPO that in previous modelling may appear distant. The *trans* model of TPO behaved in a similar manner during simulation, decreasing its maximum dimension from 100 to 82 Å. In contrast however, the CCP-like and EGF-like domains (again containing the same epitope) tended to coalesce around the reported T13 epitope residues K713-S720 on the MPO-like domain, which is also close to R225 and D707 epitope residues (Figures 7, 8, S5 & S6C). Again, the distance between Y766-Q775 and K713-S720 in the *trans* model prior to simulation is 51 Å, but approached 26 Å during simulation. For the extended model the maximum dimension reduced from 130 Å to 83 Å during the simulation. However, the CCP-like and EGF-like domains condensed in a less homogeneous manner compared to the *cis* and *trans* models, such that it was able to adopt multiple conformations relative to the MPO-like domain. Despite this, it is able to stably adopt conformations that result in all four published epitopes for the T13 autoantibody colocalising to form a continuous epitope such that they could all be plausibly engaged by a single autoantibody (Figures 7C & S6B). One such conformation selected from a stable state provided by our MD simulation suggest that all residues involved in these four epitopes are no more than 43 Å away from all other residues in the set of 39 residues previously implicated in the T13 epitope (Figures 8C & S6B). Taken together, these observations may suggest a mechanism not only by which TPO can alter its conformation to take a more compact and globular structure, but potentially explain how disparate residues involved in the IDRs may compact to form a discrete, continuous epitope.

**Figure 7.**
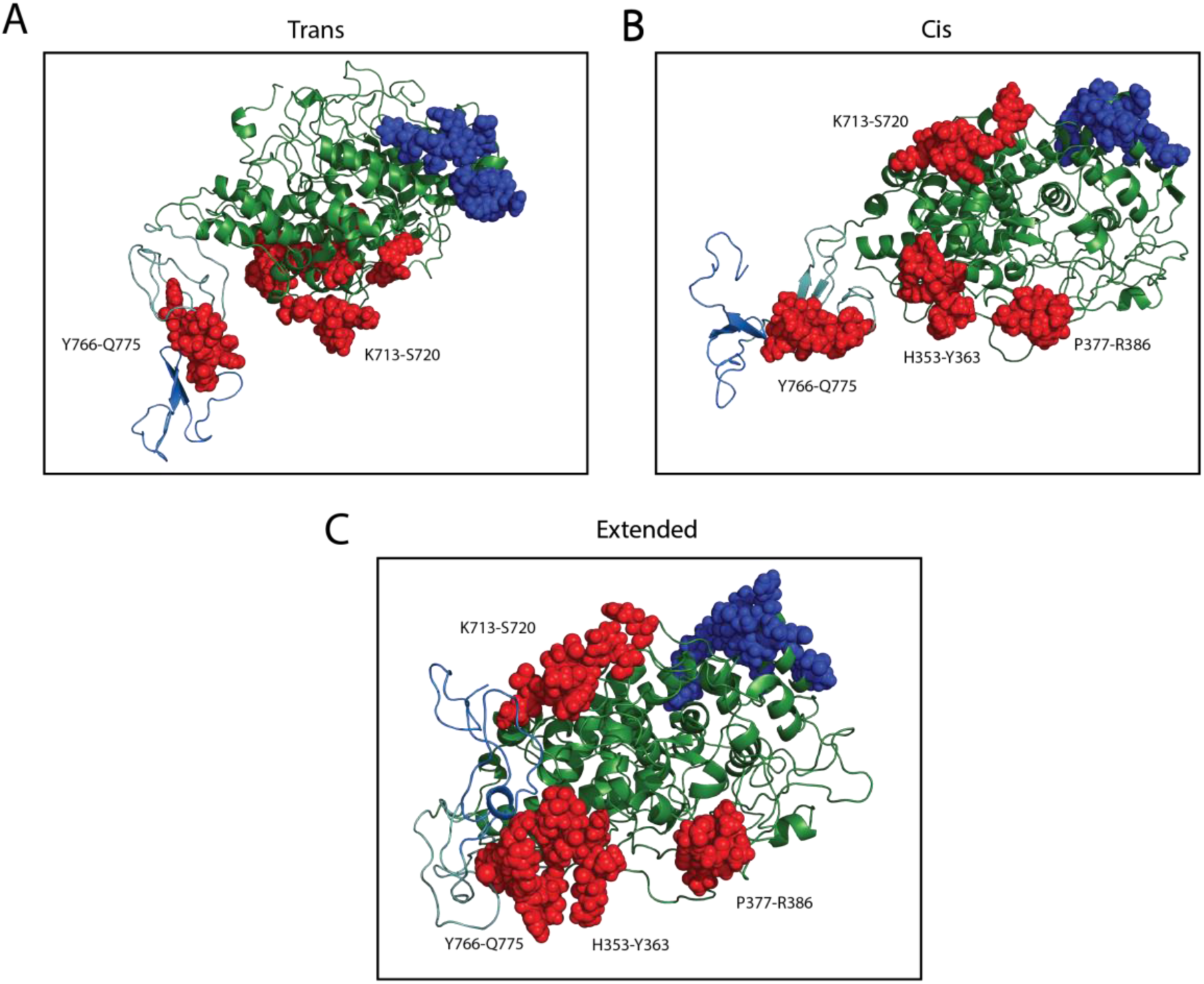
IDRs in the context of the MD simulations. A representative structure taken from the MD simulations at equilibrium for each of the (**A)** *trans*, (**B)** *cis* and (**C)** extended forms of the ΔproTPOe monomer. IDR-A residues are highlighted by red spheres, and IDR-B residues by blue spheres. The MPO-like domain, CCP-like domain and EGF-like domain are coloured in forest green, light teal and marine blue respectively (as in Figure 1). **(A)** The IDR-A epitopes of K713-S720 and Y766-Q775 are 28 Å apart from each other in this selected frame from the *trans* MD simulation. **(B)** In the *cis* model, Y766-Q775 is 21 Å from the H353-Y363 epitope and 40 Å away from the P377-R386 epitopes. **(C)** In this representation of the *extended* model during the MD simulation, all four labelled epitopes of IDR-A are no more than 43 Å away from each other. A comparison showing these structures compared with the starting TPO models is shown in Figure S6.

**Figure 8.**
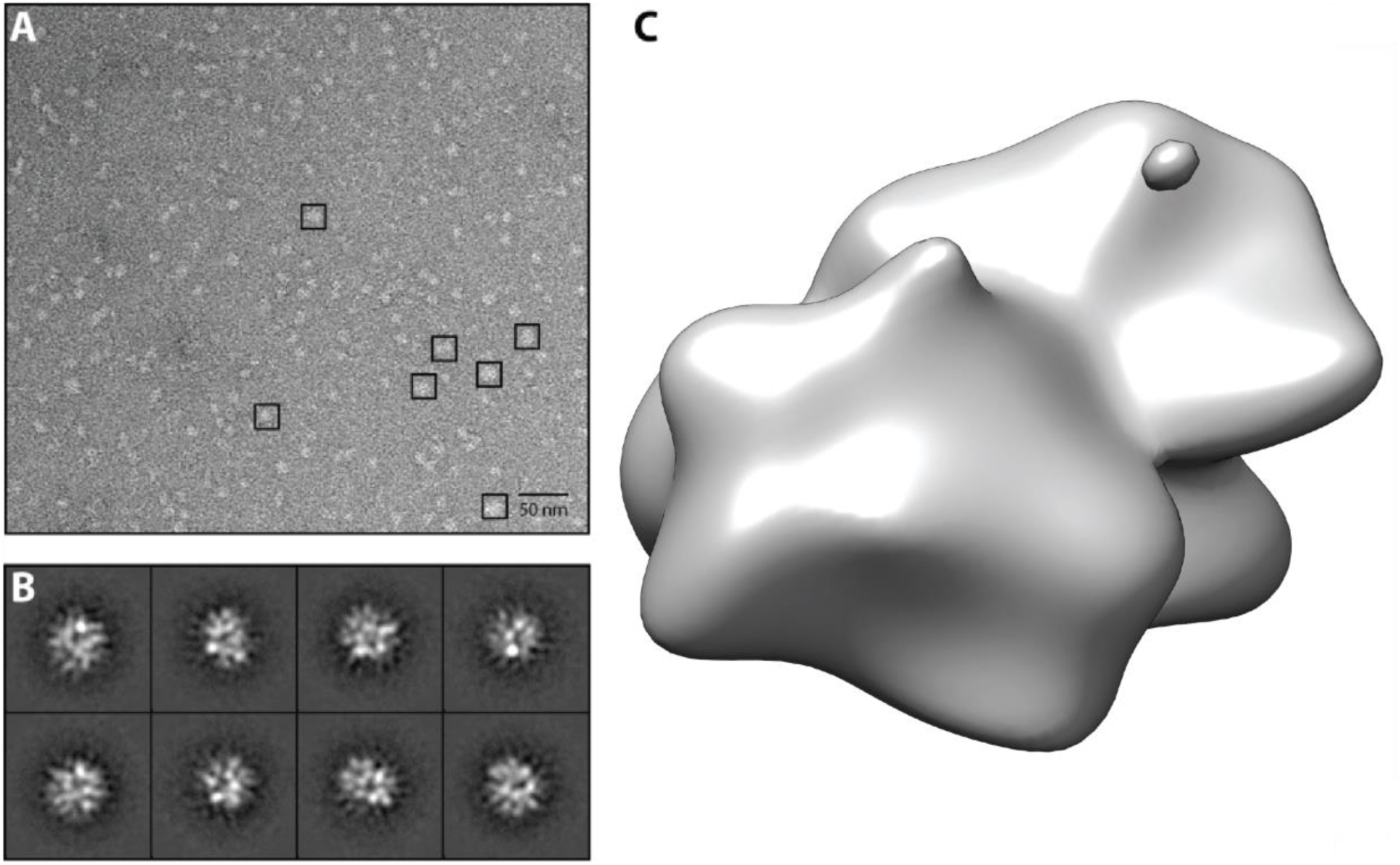
Electron micrograph of ΔproTPOe-8His in complex with TR1.9 Fab. **(A)** Representative micrograph of the ΔproTPOe-8His/TR1.9 complex collected using a FEI Tecnai Spirit T12 TEM in low-dose conditions. **(B)** 2D class averages generated from the single particles picked in (A) and used for initial 3D model generation. **(C)** 3D reconstruction of the EM volume.

### Analysis of TPO Structure Using Negative Stain Electron Microscopy

We next investigated the structure of the ΔproTPOe-8His construct in complex with TR1.9 Fab using negative stain electron microscopy. The most homogenous population of the TPO-Fab complex was selected using size-exclusion chromatography (Figure 5). The representative 2D class averages show a globular compact protein complex and initial 3D reconstruction stalled at a resolution of ∼20 Å (Figure 8). Although this limitation in resolution means that we cannot unambiguously determine the structure of a TPO-Fab complex, we can however rule out certain configurations. Docking of various models of Fab-bound monomeric and dimeric TPO into the EM envelope suggests that various configurations are feasible. Visual inspection suggests that ΔproTPOe adopts a relatively compact, rather than extended conformation, consistent with our MD analysis. For further modelling we therefore proceeded using the aforementioned compact conformations resulting from MD simulations for further analysis.

First, we sought to determine if fitting might provide information germane to the oligomerisation state of TPO. For simplicity, only the *trans* arrangements of ΔproTPOe will be presented here. There is sufficient space in the EM envelope for either a monomer or dimer of TPO, or a monomer of TPO bound to Fab. There is insufficient space however, for a dimer with two bound Fabs, and so this possibility was rejected. A Fab molecule was fitted to the envelope with the complementarity determining regions (CDRs) positioned at a distance of 4 Å from the previously published epitope of TR1.9, K713-S720. This is consistent with studies showing an antibody-epitope distance of 4.5-5 Å ^35, 36^. We also additionally selected a plausible conformation from our MD simulations of *trans* TPO that showed superior fitting to the EM volume, i.e. a folded or condensed monomer. Calculating the goodness of fit of molecules to the EM envelope as the percentage of molecules contained within the envelope, the *trans* dimer, *trans* monomer-Fab complex and condensed monomer-Fab complex contained 54, 53 and 58 percent of molecules within the volume, respectively (Figure 9). This would indicate that the condensed TPO monomer suggested by the MD simulations is the most likely approximation to the solution conformation, at least in the presence of Fab. Fitting the Fab in sequentially with TPO, i.e. not biasing the Fab orientation toward the published epitope, the fit improves to 68%. Despite this however, this fit does not place the CDRs facing in a realistic location – the CDRs face away from TPO itself. Intrinsic flexibility between the variable and constant domains of a Fab often results in ambiguity in fitting into EM data, with only the variable domains being resolved in EM density ^37–39^. Accordingly, fitting only the variable domains of the Fab in a TPO monomer-Fab complex within the envelope improves the fit to approximately 73% (Table S4).

**Figure 9.**
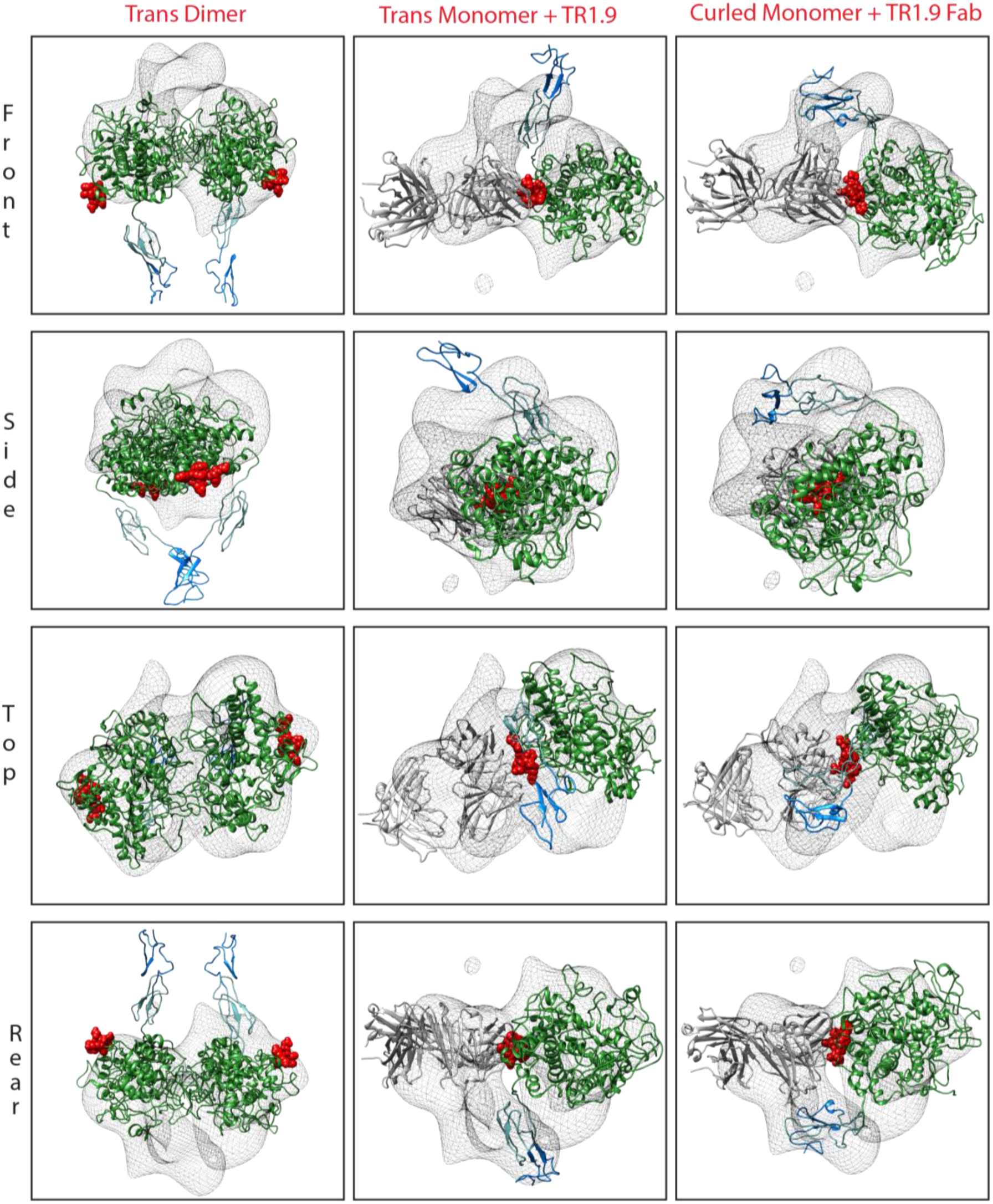
Construction of a 3D model of the ΔproTPOe-8His-TR1.9 complex via negative stain EM. 3D reconstruction of the ΔproTPOe-TR1.9 Fab complex. ΔproTPOe *trans* dimer alone, ΔproTPOe *trans* monomer with docked TR1.9 Fab, and monomer from MD simulations with docked TR1.9 Fab were fitted into the EM envelope and are marked in their respective columns. The MPO-like domain, CCP-like domain and EGF-like domain are coloured in forest green, light teal and marine blue respectively (as in Figure 1). The previously published TR1.9 epitope (K713-S720) is indicated by red spheres. Rows indicate orientation: Top and rear represent a + 90° and + 270° rotation respectively in the y axis with respect to the Front representation, while Side indicates a −90° rotation in the x axis with respect to the Front representation.

Given the orthogonal agreement within the presented data from SEC, BLItz, ELISA and AUC analysis indicating we have a high affinity binding interaction between ΔproTPOe-8His and TR1.9 Fab, it is unlikely that the Fab is not contained with the EM envelope. Unbound monomer or dimers are thus unlikely. Given that AUC analysis suggests that TR1.9 Fab binds to a TPO monomer and adopts a compact shape upon complexation that is consistent with our MD analysis (Figures 5 and 6), the EM data suggests strongly that at least in its non-membrane bound form, TR1.9 binds to TPO as a monomer.

## Discussion

Native TPO can be isolated and purified from tissue, although the availability of human thyroid tissue is a limiting factor in any preparation of TPO and rarely is enough produced for significant structural characterisation ^40^. Additionally, during maturation TPO is trimmed by proteolytic cleavage within its N-terminal propeptide region ^17, 41^. This process of proteolysis is at least in some way responsible for the low homogeneity of TPO that is purified from tissue. The ΔproTPOe-8His and ΔproTPOe-GCN4 constructs we describe improve the biochemical quality and production yield of recombinant TPO considerably, having similar functional and immunogenic properties to wild-type TPO.

The correct incorporation of the heme porphyrin ring is essential for the nascent protein’s exit from the ER and indeed for protein functionality ^16^, requiring hydrogen peroxide provided by the dual oxidase (DUOX) family of proteins present in the thyroid follicular cell membrane and are closely associated with TPO ^10, 42, 43^. Anticipating that the autocatalytic process of heme incorporation might be reduced during mammalian expression due to a lack of hydrogen peroxide, the media was supplemented with hydrogen peroxide as well as hematin, resulting in enzyme activity for ΔproTPOe-8His but not ΔproTPOe-GCN4 ^10^. Given that we demonstrate that the heme group has been incorporated, one explanation for the absence of enzyme activity for ΔproTPOe-GCN4 is possible occlusion of the active site as a result of enforced dimerization due to incorporation of the leucine zipper dimerisation motif.

Both ΔproTPOe-8His and ΔproTPOe-GCN4 constructs showed an approximately 75kDa degradation product after the initial purification, which co-eluted with the full-length protein during size-exclusion chromatography. This behavior may be due to instability or altered glycosylation, which could account for upto 25% of TPO’s 110 kDa molecular mass. There are many splice variants of TPO that have been characterised, indicating its complicated biosynthesis and the relevance of our engineered constructs in providing a consistent protein product ^40, 44^. It is now known that there is more than 10 TPO isoforms ^10^. Given the importance of glycosylation in the folding and post-translational trafficking of TPO ^9 45^, it is unlikely that non-post-translationally modified TPO would be exported. A mere 2% of TPO synthesised by cells reaches the cell surface as most is degraded by proteases or the proteasome due to partial or complete misfolding ^45^. It is possible that during the 7-day time course for expression, proteolytic activity, rather than inherent instability may have led to the observed degradation, consistent with our thermal stability data.

There exists evidence for both monomeric and dimeric forms of TPO, and no consensus agreement on its physiological oligomerization state ^17, 46–48^. Bioinformatic analysis of the peroxidase family is consistent with both monomer and dimer. LPO is the most closely related protein to TPO, with 48% sequence identity between their MPO-like domains, and is functionally monomeric ^1^. MPO, with 47% sequence identity with TPO, is an active dimer stabilized by intermolecular disulfide bonds via a conserved Cys296, which is also conserved in TPO ^48^. As such, the MPO-like domain of TPO has been modelled exclusively upon the MPO homodimer. Given however that much of the epitope mapping data suggests that autoantibodies disproportionately recognise epitopes of the MPO-like domain of TPO, and that to date TPO dimerization has not been investigated in any detail, a greater understanding of its oligomerization state would be a great benefit in understanding its autoantigenicity ^49^. Some insights may be gleaned from studies on MPO, given its common ancestry and function with TPO. MPO dimerisation occurs in the secretory pathway during proteolytic processing ^16^. In TPO biosynthesis, the propeptide is cleaved after exit from the Golgi apparatus and dimerisation is likely to occur prior to reaching the plasma membrane ^10^. As such, dimerisation may occur during the secretory pathway as in MPO, although MPO is not localized in the plasma membrane like TPO, providing additional sites at which TPO may dimerise. A small percentage (∼10%) of MPO is secreted as a fully functional monomer with exactly half the enzyme activity compared to the dimer ^16^. As such, MPO activity does not require dimerization and the functional advantages of dimerisation are not understood. It is reasonable to hypothesize therefore that TPO is functional in both monomeric and dimeric states.

AUC analysis of the ΔproTPOe-GCN4 construct is consistent with a monomer:dimer equilibrium, slightly favoring the monomer. TR1.9 Fab binds the monomer preferentially, shifting the equilibrium to 2:1, suggesting a role of the autoantibody in the monomer-dimer interplay. However, since the GCN4 leucine zipper was introduced as a dimerisation motif, any affect on dimerization by Fab-binding must be interpreted with caution. AUC analysis for ΔproTPOe-8His construct follows similar behavior in that it can exist as both monomer and dimer, with only the monomer binding to TR1.9 Fab. Interestingly, whereas the ΔproTPOe-8His construct was enzymatically active, the ΔproTPOe-GCN4 construct was not, suggesting that either the dimer interface may occlude the active site of the protein, or that GCN4 interferes with activity via another mechanism. Inspection of epitope mapping data on the *cis* model of dimeric TPO indicates that several residues that comprise immunodominant regions are buried. Furthermore, the large and diffuse nature of IDR-A and -B determinants in both *cis* and *trans* models suggest that significant conformational changes would be necessary to form a continuous epitope compatible with antibody engagement ^8^. Some residues may be surface exposed, but the dimer conformation may prevent autoantibody binding due to steric occlusion. In support of this, AUC analysis shows that the TPO-Fab complex undergoes a large conformational change to become more compact, and also that TR1.9 Fab preferentially binds the monomer, thus shifting the monomer:dimer equilibrium.

Our EM analysis provides the first ever structural glimpse regarding information about the ectodomain of TPO. Although the EM resolution is limited, it is consistent with our other data, allowing us with high confidence to eliminate a complex between Fab and a TPO dimer. Furthermore, the EM data strongly supports our SEC and AUC analysis showing that ΔproTPOe-8His is a monomer when bound to TR1.9 Fab. Modelling of TPO has suggested that membrane bound TPO is an elongated molecule with domains outstretched, however our experimental EM and AUC data suggests that at least for the TPO ectodomain, a much more globular and compact space is taken up in solution, especially when bound to its cognate antibody. This experimental data is supported by MD simulations of ΔproTPOe in its non-membrane bound form, whereby TPO is able to undergo large changes in conformation (Figure S6). Thus, MD simulations, AUC and EM provide convincing evidence that TPO is able to coalesce into a more compact, globular structure in solution. Furthermore, MD analysis reveals how the TPO monomer can change shape such that residues implicated in autoantibody binding that are disparate in space can be bought together to form a continuous epitope. However, we cannot from this data suggest definitively that this is the conformation of TPO when bound to T13 autoantibody. Instead, we argue that our data suggest that TPO may be able to change conformation in such a fashion that previously distal epitopes can coalesce into one discrete, continuous epitope. This may explain the previously conflicting epitope mapping data for some autoantibodies in IDR-A and IDR-B that contained residues that were too far from each other to be engaged by the CDRs of a single autoantibody, based on previous modelling. Despite previous attempts to resolve this issue ^14, 15^, a definitive answer, however, awaits a high-resolution structure for TPO alone and in complex with cognate autoantibodies.

All of our analysis, including MD, considers TPO in solution. We expect that the conformational, enzymatic and antigenic properties of TPO in solution are different when it is tethered to the membrane in the human thyroid. We speculate that perhaps, in the case of some TPO autoantibodies such as T13 that bind a range of spatially separate residues on TPO, when membrane bound (due to steric constraints of the membrane and other membrane bound components) the conformation required to display a continuous epitopes would be a rare event. Alternatively, this compacted conformation may become available as the result of improper trafficking to the membrane, followed by conformational change in solution. This rare conformation may perhaps be an explanation behind the observation that diseases involving TPO autoantibodies have not only a slow onset, but also explain the evidence suggesting that TPO autoantibody titre detected in patient serum does not always correlate with disease state ^50^. Perhaps only some TPO would enter this rarer, antigenic configuration, and thus lead to downstream immune effects, whilst correctly trafficked TPO or TPO in a non-antigenic configuration may remain unaffected by these autoantibodies, resulting in patients with anti-TPO antibody titre without appreciable evidence of disease. Our observation that TR1.9 Fab preferentially binds monomeric ΔproTPOe-8His may indicate that the complex (and thus pathology within the patient) appears when cryptic epitopes that are normally hidden are revealed to the immune system upon dissociation of the dimer. Our data shows that the autoantibody TR1.9 inhibits the catalytic activity of TPO constructs produced, either directly via blocking access to the active site, indirectly by favouring a conformational change that occludes the active site, or both. Although a precise understanding awaits high resolution structural analysis, this behavior suggests an intriguing connection between enzyme function and autoantigenicity that has been demonstrated for other autoantigens ^51^.

## Conclusions

Several key advances arise from this work. We designed and expressed two TPO ectodomain constructs, ΔproTPOe-8His and ΔproTPOe-GCN4, representing new reagents with which to study TPO structure, function and antigenicity. Biophysical characterisation using AUC suggests that the TPO ectodomain can exist as both a monomer and a dimer. EM and molecular dynamics analysis shows that the Fab fragment of the patient-derived TR1.9 autoantibody preferentially binds the TPO monomer, and suggests conformational change that consolidates antibody-binding residues into a continuous epitope. Taken together, these data represent the first glimpses of the structural characteristics of TPO, which thus far has resisted structural characterization by any technique. As a whole, our findings advance our understanding of TPO’s structure and function, as well as its role as a major autoantigen in autoimmune thyroid diseases.

## Acknowledgements

TR 1.9 Fab construct was kindly provided by Sandra McLachlan and Basil Rappoport. We would like to thank Dr. David Steer for help with MS analysis.

## Data Availability

The datasets generated during and/or analyzed during the current study are not publicly available but are available from the corresponding author on reasonable request.

## References

1. Williams DE, Le SN, Godlewska M, Hoke DE, Buckle AM. Thyroid Peroxidase as an Autoantigen in Hashimoto’s Disease: Structure, Function, and Antigenicity. Hormone and metabolic research = Hormon- und Stoffwechselforschung = Hormones et metabolisme. 2018;50(12):908–921.

2. Fröhlich E, Wahl R. Thyroid Autoimmunity: Role of Anti-thyroid Antibodies in Thyroid and Extra-Thyroidal Diseases. Frontiers in immunology. 2017;8:521.

3. Godlewska M, Gawel D, Buckle AM, Banga JP. Thyroid Peroxidase Revisited - What’s New? Hormone and metabolic research = Hormon- und Stoffwechselforschung = Hormones et metabolisme. 2019;51(12):765–769.

4. Silva de Morais N, Stuart J, Guan H, et al. The Impact of Hashimoto Thyroiditis on Thyroid Nodule Cytology and Risk of Thyroid Cancer. Journal of the Endocrine Society. 2019;3(4):791–800.

5. Cooper GS, Stroehla BC. The epidemiology of autoimmune diseases. Autoimmunity reviews. 2003;2(3):119–125.

6. Sarkhail P, Mehran L, Askari S, Tahmasebinejad Z, Tohidi M, Azizi F. Maternal Thyroid Function and Autoimmunity in 3 Trimesters of Pregnancy and their Offspring’s Thyroid Function. Hormone and metabolic research = Hormon- und Stoffwechselforschung = Hormones et metabolisme. 2016;48(1):20–26.

7. Seror J, Amand G, Guibourdenche J, Ceccaldi PF, Luton D. Anti-TPO antibodies diffusion through the placental barrier during pregnancy. PloS one. 2014;9(1):e84647.

8. Le SN, Porebski BT, McCoey J, et al. Modelling of Thyroid Peroxidase Reveals Insights into Its Enzyme Function and Autoantigenicity. PloS one. 2015;10(12):e0142615.

9. Godlewska M, Arczewska KD, Rudzinska M, et al. Thyroid peroxidase (TPO) expressed in thyroid and breast tissues shows similar antigenic properties. PloS one. 2017;12(6):e0179066.

10. Godlewska M, Banga PJ. Thyroid peroxidase as a dual active site enzyme: Focus on biosynthesis, hormonogenesis and thyroid disorders of autoimmunity and cancer. Biochimie. 2019;160:34–45.

11. Kimura S, Kotani T, McBride OW, et al. Human thyroid peroxidase: complete cDNA and protein sequence, chromosome mapping, and identification of two alternately spliced mRNAs. Proceedings of the National Academy of Sciences of the United States of America. 1987;84(16):5555–5559.

12. Singh AK, Singh N, Sinha M, et al. Binding modes of aromatic ligands to mammalian heme peroxidases with associated functional implications: crystal structures of lactoperoxidase complexes with acetylsalicylic acid, salicylhydroxamic acid, and benzylhydroxamic acid. The Journal of biological chemistry. 2009;284(30):20311–20318.

13. Fiedler TJ, Davey CA, Fenna RE. X-ray crystal structure and characterization of halide-binding sites of human myeloperoxidase at 1.8 A resolution. The Journal of biological chemistry. 2000;275(16):11964–11971.

14. Gardas A, Sohi MK, Sutton BJ, McGregor AM, Banga JP. Purification and crystallisation of the autoantigen thyroid peroxidase from human Graves’ thyroid tissue. Biochemical and biophysical research communications. 1997;234(2):366–370.

15. Hendry E, Taylor G, Ziemnicka K, Grennan Jones F, Furmaniak J, Rees Smith B. Recombinant human thyroid peroxidase expressed in insect cells is soluble at high concentrations and forms diffracting crystals. The Journal of endocrinology. 1999;160(3):R13–15.

16. Nauseef WM. Biosynthesis of human myeloperoxidase. Archives of biochemistry and biophysics. 2018;642:1–9.

17. Godlewska M, Gora M, Buckle AM, et al. A redundant role of human thyroid peroxidase propeptide for cellular, enzymatic, and immunological activity. Thyroid : official journal of the American Thyroid Association. 2014;24(2):371–382.

18. Chacko S, Padlan EA, Portolano S, McLachlan SM, Rapoport B. Structural studies of human autoantibodies. Crystal structure of a thyroid peroxidase autoantibody Fab. The Journal of biological chemistry. 1996;271(21):12191–12198.

19. Porebski BT, Keleher S, Hollins JJ, et al. Smoothing a rugged protein folding landscape by sequence-based redesign. Scientific Reports. 2016;6:33958.

20. Pettersen EF, Goddard TD, Huang CC, et al. UCSF Chimera--a visualization system for exploratory research and analysis. J Comput Chem. 2004;25(13):1605–1612.

21. Dolinsky TJ, Nielsen JE, McCammon JA, Baker NA. PDB2PQR: an automated pipeline for the setup of Poisson–Boltzmann electrostatics calculations. Nucleic Acids Research. 2004;32(suppl_2):W665–W667.

22. Søndergaard CR, Olsson MH, Rostkowski M, Jensen JH. Improved treatment of ligands and coupling effects in empirical calculation and rationalization of p K a values. Journal of chemical theory and computation. 2011;7(7):2284–2295.

23. Jorgensen WL, Chandrasekhar J, Madura JD, Impey RW, Klein ML. Comparison of simple potential functions for simulating liquid water. The Journal of chemical physics. 1983;79(2):926–935.

24. Joung IS, Cheatham III TE. Determination of alkali and halide monovalent ion parameters for use in explicitly solvated biomolecular simulations. The journal of physical chemistry B. 2008;112(30):9020–9041.

25. Maier JA, Martinez C, Kasavajhala K, Wickstrom L, Hauser KE, Simmerling C. ff14SB: improving the accuracy of protein side chain and backbone parameters from ff99SB. Journal of chemical theory and computation. 2015;11(8):3696–3713.

26. Li P, Roberts BP, Chakravorty DK, Merz Jr KM. Rational design of particle mesh Ewald compatible Lennard-Jones parameters for+ 2 metal cations in explicit solvent. Journal of chemical theory and computation. 2013;9(6):2733–2748.

27. Berendsen HJ, Postma Jv, van Gunsteren WF, DiNola A, Haak JR. Molecular dynamics with coupling to an external bath. The Journal of chemical physics. 1984;81(8):3684–3690.

28. Phillips JC, Braun R, Wang W, et al. Scalable molecular dynamics with NAMD. Journal of computational chemistry. 2005;26(16):1781–1802.

29. Humphrey W, Dalke A, Schulten K. VMD: visual molecular dynamics. Journal of molecular graphics. 1996;14(1):33–38.

30. McGibbon RT, Beauchamp KA, Harrigan MP, et al. MDTraj: a modern open library for the analysis of molecular dynamics trajectories. Biophysical journal. 2015;109(8):1528–1532.

31. Bresson D, Cerutti M, Devauchelle G, et al. Localization of the discontinuous immunodominant region recognized by human anti-thyroperoxidase autoantibodies in autoimmune thyroid diseases. The Journal of biological chemistry. 2003;278(11):9560–9569.

32. Bresson D, Pugniere M, Roquet F, et al. Directed mutagenesis in region 713-720 of human thyroperoxidase assigns 713KFPED717 residues as being involved in the B domain of the discontinuous immunodominant region recognized by human autoantibodies. The Journal of biological chemistry. 2004;279(37):39058–39067.

33. Rebuffat SA, Bresson D, Nguyen B, Peraldi-Roux S. The key residues in the immunodominant region 353-363 of human thyroid peroxidase were identified. International immunology. 2006;18(7):1091–1099.

34. Estienne V, Duthoit C, Blanchin S, et al. Analysis of a conformational B cell epitope of human thyroid peroxidase: identification of a tyrosine residue at a strategic location for immunodominance. International immunology. 2002;14(4):359–366.

35. Ramaraj T, Angel T, Dratz EA, Jesaitis AJ, Mumey B. Antigen-antibody interface properties: composition, residue interactions, and features of 53 non-redundant structures. Biochimica et biophysica acta. 2012;1824(3):520–532.

36. Bordo D, Argos P. Evolution of protein cores. Constraints in point mutations as observed in globin tertiary structures. Journal of molecular biology. 1990;211(4):975–988.

37. Jiang Q-X, Wang D-N, MacKinnon R. Electron microscopic analysis of KvAP voltage-dependent K+ channels in an open conformation. Nature. 2004;430(7001):806–810.

38. Conway JF, Watts NR, Belnap DM, et al. Characterization of a Conformational Epitope on Hepatitis B Virus Core Antigen and Quasiequivalent Variations in Antibody Binding. Journal of Virology. 2003;77(11):6466–6473.

39. Wu S, Avila-Sakar A, Kim J, et al. Fabs enable single particle cryoEM studies of small proteins. Structure (London, England : 1993). 2012;20(4):582–592.

40. Gardas A, Lewartowska A, Pasieka Z, Sutton BJ, McGregor AM, Banga JP. Human Thyroid Peroxidase (TPO) Isoforms, TPO-1 and TPO-2: Analysis of Protein Expression in Graves’ Thyroid Tissue1. The Journal of Clinical Endocrinology & Metabolism. 1997;82(11):3752–3757.

41. Godlewska M, Krasuska W, Czarnocka B. Biochemical properties of thyroid peroxidase (TPO) expressed in human breast and mammary-derived cell lines. PloS one. 2018;13(3):e0193624.

42. de Carvalho DP, Lima de Souza EC, Fortunato RS, et al. Functional Consequences of Dual Oxidase-Thyroperoxidase Interaction at the Plasma Membrane. The Journal of Clinical Endocrinology & Metabolism. 2010;95(12):5403–5411.

43. Carvalho DP, Dupuy C. Role of the NADPH Oxidases DUOX and NOX4 in Thyroid Oxidative Stress. European thyroid journal. 2013;2(3):160–167.

44. Elisei R, Vassart G, Ludgate M. Demonstration of the existence of the alternatively spliced form of thyroid peroxidase in normal thyroid. The Journal of clinical endocrinology and metabolism. 1991;72(3):700–702.

45. Fayadat L, Niccoli-Sire P, Lanet J, Franc JL. Role of heme in intracellular trafficking of thyroperoxidase and involvement of H2O2 generated at the apical surface of thyroid cells in autocatalytic covalent heme binding. The Journal of biological chemistry. 1999;274(15):10533–10538.

46. de Vijlder JJM, Bikker H. Biochemistry and Physiology of Thyroid Peroxidase. 2000; Berlin, Heidelberg.

47. Baker JR, Arscott P, Johnson J. An analysis of the structure and antigenicity of different forms of human thyroid peroxidase. Thyroid : official journal of the American Thyroid Association. 1994;4(2):173–178.

48. McDonald DO, Pearce SH. Thyroid peroxidase forms thionamide-sensitive homodimers: relevance for immunomodulation of thyroid autoimmunity. Journal of molecular medicine (Berlin, Germany). 2009;87(10):971–980.

49. Godlewska M, Czarnocka B, Gora M. Localization of key amino acid residues in the dominant conformational epitopes on thyroid peroxidase recognized by mouse monoclonal antibodies. Autoimmunity. 2012;45(6):476–484.

50. McLachlan SM, Rapoport B. Thyroid peroxidase autoantibody epitopes revisited*. Clinical endocrinology. 2008;69(4):526–527.

51. Kass I, Hoke DE, Costa MGS, et al. Cofactor-dependent conformational heterogeneity of GAD65 and its role in autoimmunity and neurotransmitter homeostasis. Proceedings of the National Academy of Sciences. 2014;111(25):E2524–E2529.

52. Wiles AP, Shaw G, Bright J, Perczel A, Campbell ID, Barlow PN. NMR studies of a viral protein that mimics the regulators of complement activation. Journal of molecular biology. 1997;272(2):253–265.

53. Downing AK, Knott V, Werner JM, Cardy CM, Campbell ID, Handford PA. Solution structure of a pair of calcium-binding epidermal growth factor-like domains: implications for the Marfan syndrome and other genetic disorders. Cell. 1996;85(4):597–605.

