## Supporting Information for "Structural studies of thyroid peroxidase show the monomer interacting with autoantibodies in thyroid autoimmune disease"

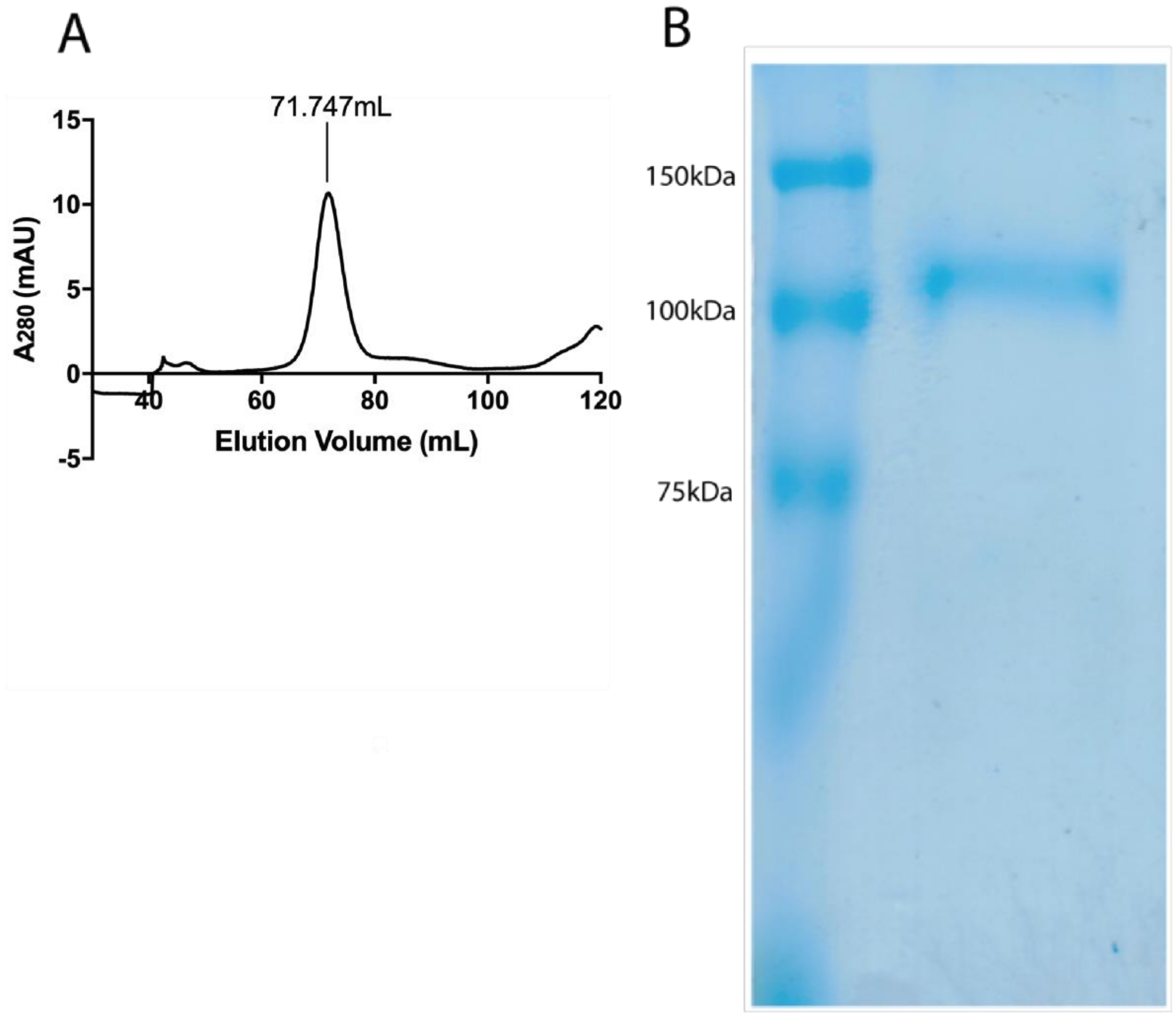


**Figure S1 - Purification of the TPO construct ΔproTPOe-GCN4 (A)** A chromatogram from a Superdex S200 16/60 column, showing ΔproTPOe-GCN4 eluting as a single major peak at 71.7mL, consistent with a 110 kDa protein. No other major large species appears to be present. **(B)** Reducing SDS-PAGE analysis of purified ΔproTPOe-GCN4 shows a major band at ~110 kDa.

1 MRALAVLSVT LVMACTEAFF PFISRGKELL WGKPEESRVS SVLEESKRLV

51 DTAMYATMQR NLKKRGILSP AQLLSFSKLP EPTSGVIARA AEIMETSIQA

101 MKRKVNLKTQ QSQHPTDALS EDLLSIIANM SGCLPYMLPP KCPNTCLANK

151 **YRPITGACNN** **R**DHPR**WGASN** **TALARWLPPV** **YEDGFSQPRG** **WNPGFLYNGF**

201 **PLPPVR**EVTR **HVIQVSNEVV** **TDDDRYSDLL** **MAWGQYIDHD** **IAFTPQSTSK**

251 AAFGGGADCQ MTCENQNPCF PIQLPEEAR**P** **AAGTACLPFY** **R**SSAACGTGD

301 QGALFGNLST ANPR**QQMNGL** **TSFLDASTVY** **GSSPALER**QL RNW**TSAEGLL**

351 **R**VHARLRDSG R**AYLPFVPPR** **APAACAPEPG** **IPGETRGPCF** **LAGDGRASEV**

401 **PSLTALHTLW** **LR**EHNR**LAAA** **LKALNAHWSA** **DAVYQEARKV** **VGALHQIITL**

451 **R**DYIPR**ILGP** **EAFQQYVGPY** **EGYDSTANPT** **VSNVFSTAAF** **RFGHATIHPL**

501 **VRRLDASFQE** **HPDLPGLWLH** **QAFFSPWTLL** **RGGGLDPLIR** GLLAR**PAKLQ**

551 **VQDQLMNEEL** **TERLFVLSNS** **STLDLASINL** **QRGRDHGLPG** **YNEWREFCGL**

601 **PRLETPADLS** **TAIASRSVAD** **KILDLYKHPD** **NIDVWLGGLA** **ENFLPR**AR**TG**

651 **PLFACLIGK**Q MKAL**RDGDWF** **WWENSHVFTD** **AQR**R**ELEKHS** **LSRVICDNTG**

701 **LTRVPMDAFQ** **VGK**FPEDFES CDSITGMNLE AWR**ETFPQDD** **KCGFPESVEN**

751 **GDFVHCEESG** **R**R**VLVYSCRH** **GYELQGR**EQL TCTQEGWDFQ PPLCK**DVNEC**

801 **ADGAHPPCHA** **SAR**CRNTK**GG** **FQCLCADPYE** **LGDDGR**TCVD SGRLPRRMKQ

851 LEDKVEELLS KNYHLENEVA RLKKLVGERG TGSHHHHHH H

**Figure S2 – Mass spectrometry analysis of ΔproTPOe-GCN4**. Sequence coverage was reported as 70% with a protein score of 19294, making ΔproTPOe-GCN4 the most abundant species in the sample. Full length TPOe-GCN4 is in black lettering, with detected peptides highlighted in red. Note that residues 1 through 108 comprise the signal peptide and propeptide that are not incorporated into full length ΔproTPOe-GCN4, though are included here to demonstrate their successful non-inclusion in our construct.

```
  1 MRALAVLSVT LVMACTEAFF PFISRGKELL WGKPEESRVS SVLEESKRLV
 51 DTAMYATMQR NLKKRGILSP AQLLSFSKLP EPTSGVIARA AEIMETSIQA
101 MKRKVNLKTQ QSQHPTDALS EDLLSIIANM SGCLPYMLPP KCPNTCLANK
151 YRPITGACNN RDHPRWGASN TALARWLPPV YEDGFSQPRG WNPGFLYNGF
201 PLPPVREVTR HVIQVSNEVV TDDDRYSDLL MAWGQYIDHD IAFTPQSTSK
251 AAFGGGADCQ MTCENQNPCF PIQLPEEARP AAGTACLPFY RSSAACGTGD
301 QGALFGNLST ANPRQQMNGL TSFLDASTVY GSSPALERQL RNWTSAEGLL
351 RVHARLRDSG RAYLPFVPPR APAACAPEPG IPGETRGPCF LAGDGRASEV
401 PSLTALHTLW LREHNRLAAA LKALNAHWSA DAVYQEARKV VGALHQIITL
451 RDYIPRILGP EAFQQYVGPY EGYDSTANPT VSNVFSTAAF RFGHATIHPL
501 VRRLDASFQE HPDLPGLWLH QAFFSPWTLL RGGGLDPLIR GLLARPAKLQ
551 VQDQLMNEEL TERLFVLSNS STLDLASINL QRGRDHGLPG YNEWREFCGL
601 PRLETPADLS TAIASRSVAD KILDLYKHPD NIDVWLGGLA ENFLPRARTG
651 PLFACLIGKQ MKALRDGDWF WWENSHVFTD AQRRELEKHS LSRVICDNTG
701 LTRVPMDAFQ VGKFPEDFES CDSITGMNLE AWRETFPQDD KCGFPESVEN
751 GDFVHCEESG RRVLVYSCRH GYELQGREQL TCTQEGWDFQ PPLCKDVNEC
801 ADGAHPPCHA SARCRNTKGG FQCLCADPYE LGDDGRTCVD SGRLPRRMKQ
851 LEDKVEELLS KNYHLENEVA RLKKLVGERG TGSHHHHHHH H
```

**Figure S3 – Mass spectrometry analysis of suspected degraded ΔproTPOe-GCN4 fragment**. Sequence coverage was reported as 63% with a protein score of 8710, making a degraded form of ΔproTPOe-GCN4 the most abundant species in the sample. Full length ΔproTPOe-GCN4 is in black lettering, with detected peptides highlighted in red. Note that residues 1 through 108 comprise the signal peptide and propeptide that are not incorporated into full length ΔproTPOe-GCN4, though are included here to demonstrate their successful non-inclusion in our construct.

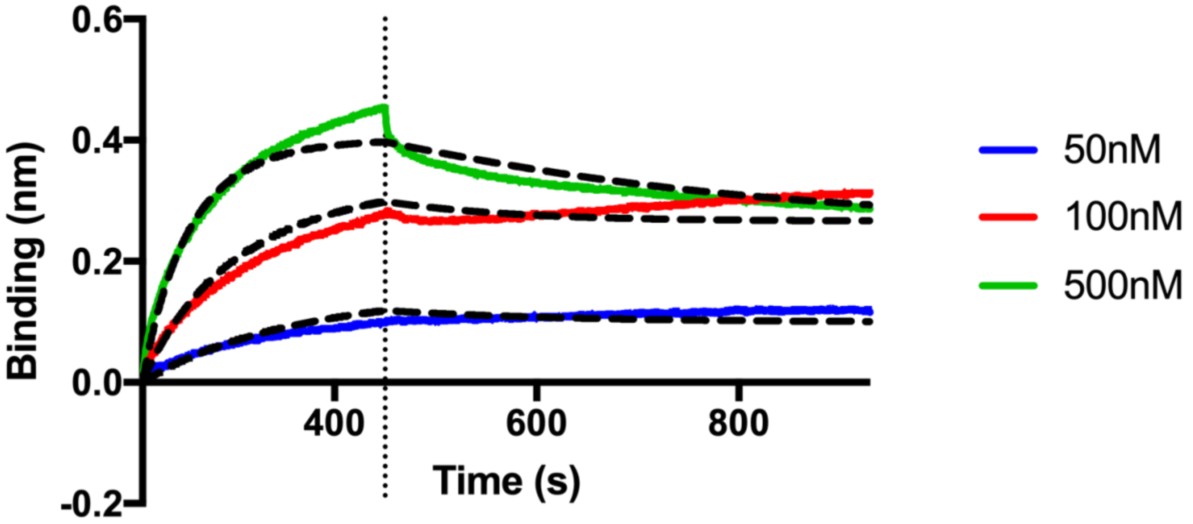


**Figure S4 – Bio-layer interferometry (BLI) sensorgram data of ΔproTPOe-8His binding to Fab**. Sensorgram curves according to a TR1.9 Fab concentration range of between 0 and 500 nM. ΔproTPOe-8His is immobilised on the biosensor surface. The data has been normalised against a blank run of buffer (1x PBS, pH 7.4). Vertical line at 450 s represents the end of the association phase. Dotted lines in black represent the fit calculated using a 1:1 binding model with global fitting within the BLItz Pro software. $R_2$ values for the calculated fit were reported as 0.97. $K_D$ was calculated as 20 nM.

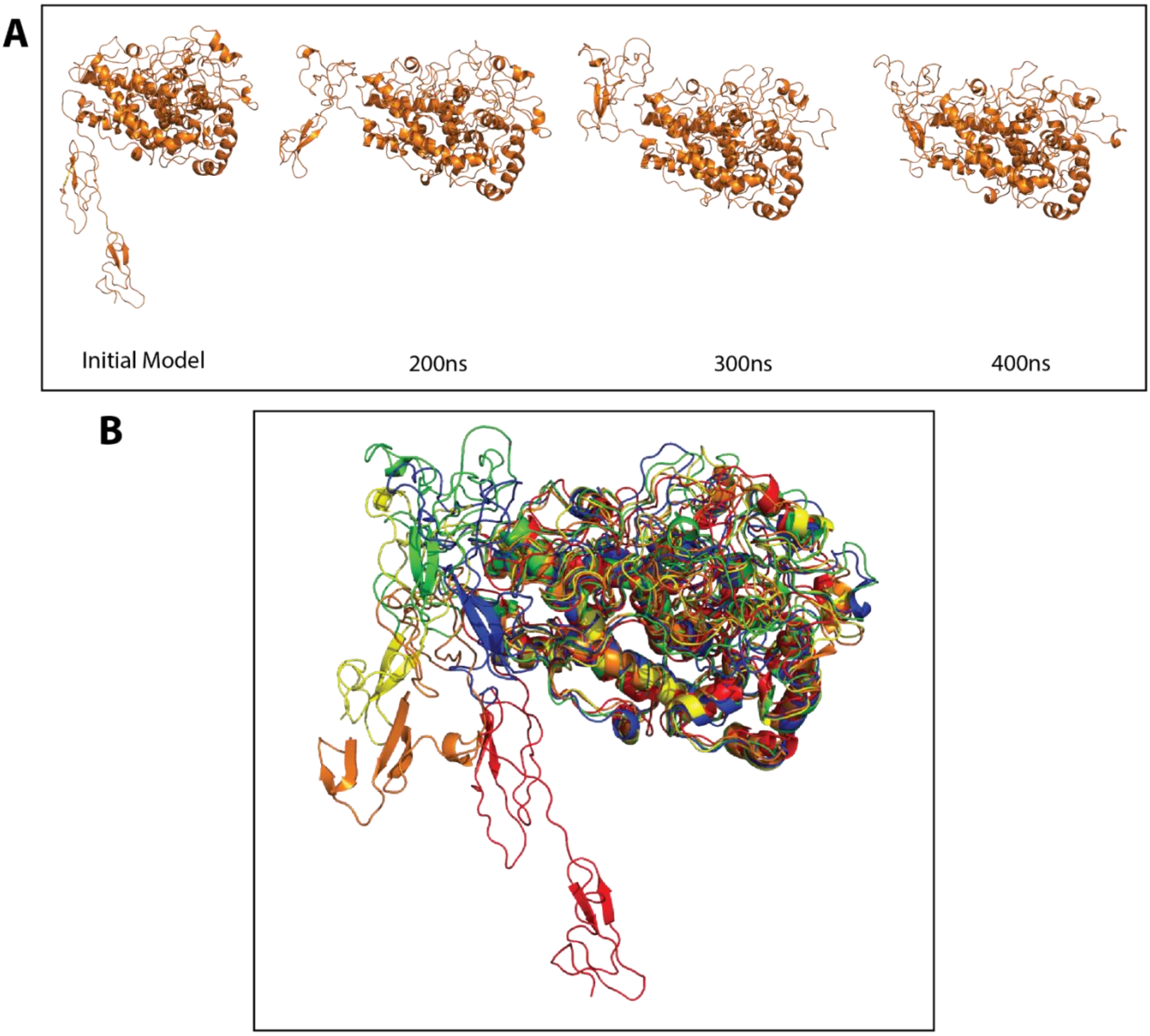


**Figure S5 – Snapshots from the trans ΔproTPOe MD trajectory show TPO changing conformation from extended to more compact structure**. Snapshots from the trans ΔproTPOe MD trajectory as presented in Figure 6. **(A)** Representation of the starting model from Le and co-workers [1], as well as *trans* ΔproTPOe after 200, 300 and 400 ns of simulation. **(B)** Structural superpositions of the above snapshots with the starting trans ΔproTPOe model in red. Orange indicates *trans* ΔproTPOe after 100 ns of simulation, yellow after 200 ns, green after 300 ns and blue after 400 ns.

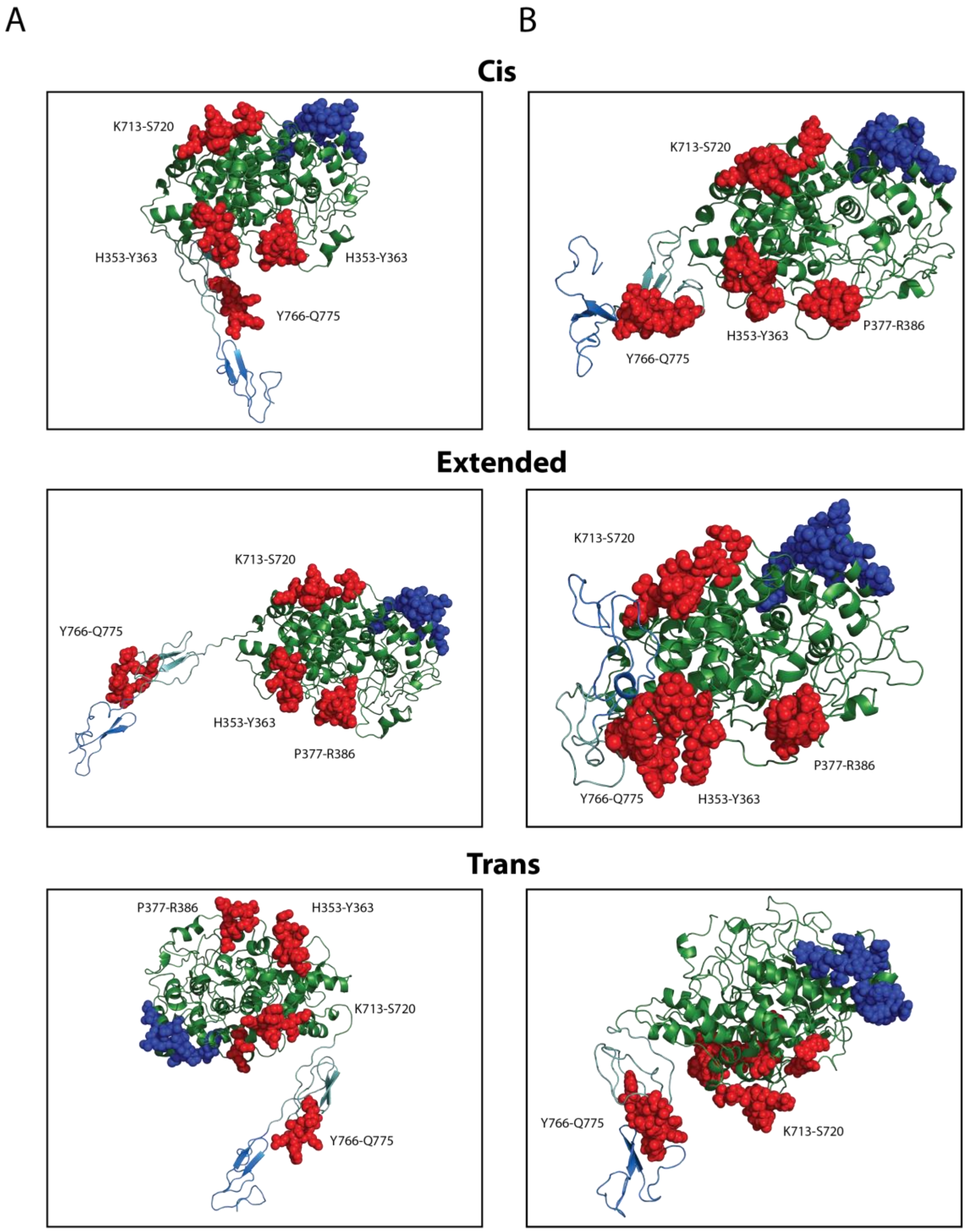


**Figure S6 – IDRs in context of the MD simulations with starting structures. (A)** TPO models (simulation starting structures) with IDRs highlighted. (**B)** Representative structures taken from the MD simulations for each of the *trans*, *cis* and *extended* forms of the ΔproTPOe monomer, as in Figure 7. IDR-A residues are highlighted by red spheres, and IDR-B residues by blue spheres. The MPO-like domain, CCP-like domain and EGF-like domain are coloured in forest green, light teal and marine blue respectively (as in Figure 1).

**Table S1** – Published residues involved in IDRs of TPO

| Antibody Involved | Number of Reported Epitopes | Epitopes | Study |
|---|---|---|---|
| **IDR-A** | | | |
| T13 | 4 | H353-Y363, P377-R386, K713-S720, Y766-Q775 | 2-6 |
| ICA1 | 1 | H353-Y363 | 2,3 |
| TR1.9 | 2 | K713, K713-S720 | 2,4,7 |
| 126TO10 | 3 | R225, R646, D707 | 8,9 |
| 126TP1 | 3 | R225, R646, D707 | 8,9 |
| 126TP7 | 1 | R225 | 9 |
| **IDR-B** | | | |
| 126TP5 | 5 | D620, D624, K627, D630, F597-E604 | 8-10 |
| 126TP14 | 5 | D620, D624, K627, D630, F597-E604 | 8-10 |
| 131TP7 | 1 | K627 | 9 |
| SP1.4 | 1 | F597-E604 | 10 |
| TR1.8 | 1 | T611-V618 | 10 |
| WR1.7 | 1 | F597-E604 | 10 |

The epitopes that have been identified as making up the immunodominant regions (IDRs) of TPO, named IDR-A and IDR-B.

**Table S2 – Melting point data of ΔproTPOe-8His in different buffer conditions**

| **Buffer** | **pH** | **$T_m$ (**°C) |
|---|---|---|
| 50 mM HEPES, 250 mM NaCl | 8.0 | 52.3 |
| 5 0mM HEPES, 250 mM NaCl | 7.0 | 55.2 |
| 50 mM Sodium Phosphate, 250 mM NaCl | 6.0 | 54.7 |
| 50 mM Sodium Acetate, 250 mM NaCl | 5.5 | 53.9 |
| 50 mM Glycine, 250 mM NaCl | 4.0 | 53.6 |

**Table S3** – Theoretical and calculated Stokes Radii ($R_s$) of TPO

| **Reference Dataset** | **Equation** | **Stokes Radius (Å)** | |
|---|---|---|---|
| | | **Monomer** | **Dimer** |
| Globular Folded Proteins | $R_s = (4.75)N_{0.29}$ | 32.52 | 39.75 |
| Denatured Unfolded Proteins | $R_s = (2.21)N_{0.57}$ | 96.93 | 143.89 |
| Analytical SEC of TPO with no TM domain | | 51.31 | |
| AUC of ΔproTPOe-GCN4 | | 75.7 | 91.4 |
| AUC of ΔproTPOe-8His | | 64.9 | 77.1 |
| AUC of ΔproTPOe-GCN4 with TR1.9 | | 52.7 | 66.4 |
| AUC of ΔproTPOe-8His with TR1.9 | | 57.4 | N/A |

AUC, analytical ultracentrifugation; SEC, size exclusion chromatography; TM, transmembrane domain.

**Table S4 – Model Fit Percentages**

| Model | Percentage of Molecules within the EM Map |
|---|---|
| Trans Monomer | 73 |
| Cis Monomer | 72 |
| Trans Dimer | 54 |
| Cis Dimer | 59 |
| Trans Monomer with Fab | 53 |
| Cis Monomer with Fab | 55 |
| Curled Monomer with Fab | 58 |
| Curled Monomer with scFv format of TR1.9 | 73 |
| Trans Monomer with Fab sequentially fit* | 68 |
| Cis Monomer with Fab sequentially fit* | 70 |
| Trans Dimer with Fab | 33 |
| Cis Dimer with Fab | 37 |

Fit percentages of various TPO models within the electron microscopy (EM) map. Asterisks (*) indicates configurations where TR 1.9 Fab was fitted into the available space in the envelope without regard to its epitope's location, rather than in a realistic orientation in which the complementarity determining regions (CDR) face the published epitope of K713-S720.

### References


1. Le SN, Porebski BT, McCoey J, et al. Modelling of Thyroid Peroxidase Reveals Insights into Its Enzyme Function and Autoantigenicity. *PloS one.* 2015;10(12):e0142615.
2. Bresson D, Cerutti M, Devauchelle G, et al. Localization of the discontinuous immunodominant region recognized by human anti-thyroperoxidase autoantibodies in autoimmune thyroid diseases. *The Journal of biological chemistry.* 2003;278(11):9560-9569.
3. Rebuffat SA, Bresson D, Nguyen B, Peraldi-Roux S. The key residues in the immunodominant region 353-363 of human thyroid peroxidase were identified. *International immunology.* 2006;18(7):1091-1099.
4. Bresson D, Pugniere M, Roquet F, et al. Directed mutagenesis in region 713-720 of human thyroperoxidase assigns 713KFPED717 residues as being involved in the B domain of the discontinuous immunodominant region recognized by human autoantibodies. *The Journal of biological chemistry.* 2004;279(37):39058-39067.
5. Williams DE, Le SN, Godlewska M, Hoke DE, Buckle AM. Thyroid Peroxidase as an Autoantigen in Hashimoto's Disease: Structure, Function, and Antigenicity. *Hormone and metabolic research = Hormon- und Stoffwechselforschung = Hormones et metabolisme.* 2018;50(12):908-921.
6. Estienne V, Duthoit C, Blanchin S, et al. Analysis of a conformational B cell epitope of human thyroid peroxidase: identification of a tyrosine residue at a strategic location for immunodominance. *International immunology.* 2002;14(4):359-366.
7. Guo J, Yan XM, McLachlan SM, Rapoport B. Search for the autoantibody immunodominant region on thyroid peroxidase: epitopic footprinting with a human monoclonal autoantibody locates a facet on the native antigen containing a highly conformational epitope. *Journal of immunology (Baltimore, Md : 1950).* 2001;166(2):1327-1333.
8. Dubska M, Banga JP, Plochocka D, et al. Structural insights into autoreactive determinants in thyroid peroxidase composed of discontinuous and multiple key contact amino acid residues contributing to epitopes recognized by patients' autoantibodies. *Endocrinology.* 2006;147(12):5995-6003.
9. Gora M, Gardas A, Watson PF, et al. Key residues contributing to dominant conformational autoantigenic epitopes on thyroid peroxidase identified by mutagenesis. *Biochemical and biophysical research communications.* 2004;320(3):795-801.
10. Bresson D, Rebuffat SA, Nguyen B, Banga JP, Gardas A, Peraldi-Roux S. New Insights into the Conformational Dominant Epitopes on Thyroid Peroxidase Recognized by Human Autoantibodies. *Endocrinology.* 2005;146(6):2834-2844.
